# Transcriptomes and metabolism define mouse and human MAIT cell heterogeneity

**DOI:** 10.1101/2021.12.20.473182

**Authors:** Shilpi Chandra, Gabriel Ascui, Thomas Riffelmacher, Ashu Chawla, Ciro Ramirez-Suastegui, Viankail Cedillo Castelan, Gregory Seumois, Hayley Simon, Mallory Paynich Murray, Goo-Young Seo, Ashmitaa Logandha Ramamoorthy Premlal, Greet Verstichel, Yingcong Li, Chia-Hao Lin, Jason Greenbaum, John Lamberti, Raghav Murthy, John Nigro, Hilde Cheroutre, Christian H. Ottensmeier, Stephen M. Hedrick, Li-Fan Lu, Pandurangan Vijayanand, Mitchell Kronenberg

## Abstract

Mucosal-associated invariant T (MAIT) cells are a subpopulation of T lymphocytes that respond to microbial metabolites. We performed single-cell RNA sequencing and metabolic analyses of MAIT cell subsets in thymus and peripheral tissues from mice and humans to define the heterogeneity and developmental pathway of these innate-like lymphocytes. We show that the predominant mouse subset, which produces IL-17 (MAIT17), and the subset that produces IFNγ (MAIT1), have greatly different transcriptomes and metabolic states in the thymus and periphery. A splenic MAIT subset has a transcriptome similar to circulating lymphocytes, and in mice these also are found in recent thymic emigrants, suggesting partially mature cells emigrate from the thymus. Human MAIT cells are predominantly MAIT1 cells, but have a different metabolism from their mouse counterparts with increased fatty acid uptake and storage. Although mouse and human subsets are similar in thymus, in the periphery they diverge, likely reflecting environmental influences.

## Introduction

Mucosal-associated invariant T (MAIT) cells are found in humans, mice and many other mammals^1^. They recognize MR1, a non-polymorphic major histocompatibility complex (MHC)-class I-like protein that binds to 5-(2-oxopropylideneamino)-6-D-ribitylaminouracil (5-OP-RU) and other riboflavin-derived metabolites produced by bacteria and yeast ^2, 3, 4, 5, 6, 7^. MAIT cells are abundant in humans, but relatively rare in laboratory mice^8, 9, 10^. In humans, MAIT cells have a restricted αβ T cell receptor (TCR), in which the TCRVα chain comprises a canonical Vα7.2-Jα33 (TRAV1-2-TRAJ33) rearrangement, paired with a limited number of TCRβ chains. The mouse MAIT cell TCR is made up predominantly of a homologous Vα19-Jα33 (TRAV1-TRAJ33) TCRα chain associated with a limited set of Vβ segments ^3, 4, 11^. Activated MAIT cells proliferate, and rapidly secrete pro-inflammatory cytokines, as well as cytotoxic effector molecules such as perforin and granzymes^12, 13^. These properties suggest that MAIT cells function as first responders to microbial infections, while in some cases, contributing to abnormal inflammatory reactions^14, 15^.

Like other T cell populations, MAIT cells originate in the thymus, but their positive selection is dependent on double positive thymocytes, similar to invariant Natural Killer T (iNKT) cells^16, 17^. A stepwise development based on the surface expression of CD24 and CD44 has been described previously for MAIT cells in mouse thymus^9, 10^. Similarly, for humans, thymus stages are defined based on expression of CD27 and CD161^9, 16^. Some thymic MAIT cells exhibit effector functions typical of differentiated peripheral T cells, a property found in other innate-like T cells including iNKT cells and γδ T cells^18, 19^.

MAIT cell heterogeneity has been demonstrated by several recent reports^10, 20, 21, 22, 23^. The predominant mouse MAIT cell subset in the thymus and elsewhere is characterized by the expression of RORγT and other surface markers, as well as IL-17 secretion after activation, and therefore they are considered to be MAIT17 cells, analogous to CD4^+^ Th17 cells. A T-bet-expressing MAIT1 population that secretes IFNγ also has been characterized^8, 9, 24, 25^. In humans, most MAIT cells fit into the MAIT1 category, but a minority of MAIT cells capable of producing IL-17 have been found in tissues^26, 27^. In contrast to conventional T cells, MAIT cells may encounter their natural antigen during thymic differentiation as metabolites from riboflavin-synthesizing bacteria enter the thymus^28^. As a result, MAIT cells are agonist-selected and appear antigen-experienced, and in addition to their immediate effector functions, they also express memory markers^29^. Memory-like vs effector-like states in conventional CD4^+^ and CD8^+^ T cells are controlled by mutually exclusive metabolic states^30, 31^, and thus the question remains as to which programs MAIT cells adopt at steady-state. As natural effector cells, we might expect that the metabolic state of MAIT cells would be different from naïve CD4 and CD8 T cells. Therefore, here we have taken an unbiased approach to characterize MAIT cell heterogeneity in mouse and human cells from different organs by analyzing transcriptomes and metabolic parameters. Our data confirm the acquisition of effector function and also a different metabolic state, even in the thymus. Furthermore, the data reveal additional MAIT cell subsets, including circulatory MAIT cells, that include precursors of more mature cells. Additionally, we show that despite the conservation of their specificities, the peripheral subsets of human and mouse MAIT cells defined here are not highly similar, and the status of mouse MAIT cells is subject to influences from the environment.

## Results

### Transcriptomic and phenotypic heterogeneity of mouse MAIT cells

To characterize mouse MAIT cells, we sorted 5-OP-RU loaded mouse MR1 tetramer^+^ cells (**Extended Data Fig. 1A**) from thymus, lung, liver and spleen from C57BL/6J mice and subjected them to single-cell RNA sequencing (scRNA-seq). We used the R package Seurat to perform a dimensional reduction and the Louvain algorithm to cluster cells. scRNA-seq of mouse MAIT cells revealed 11 clusters (**Fig. 1A and Supplementary Table 1**). Of note, the composition of the clusters was not evenly distributed across the four tissue sources **(Fig 1B and Extended Data 1B**): Cluster 6 was purely thymus-derived, clusters 3 and 7 were almost entirely from the lung, and clusters 1 and 8 had the highest proportion of cells from the liver, with relatively little representation of lung MAIT cells (**Fig. 1B, Extended Data Fig. 1B**).

**Figure 1:**
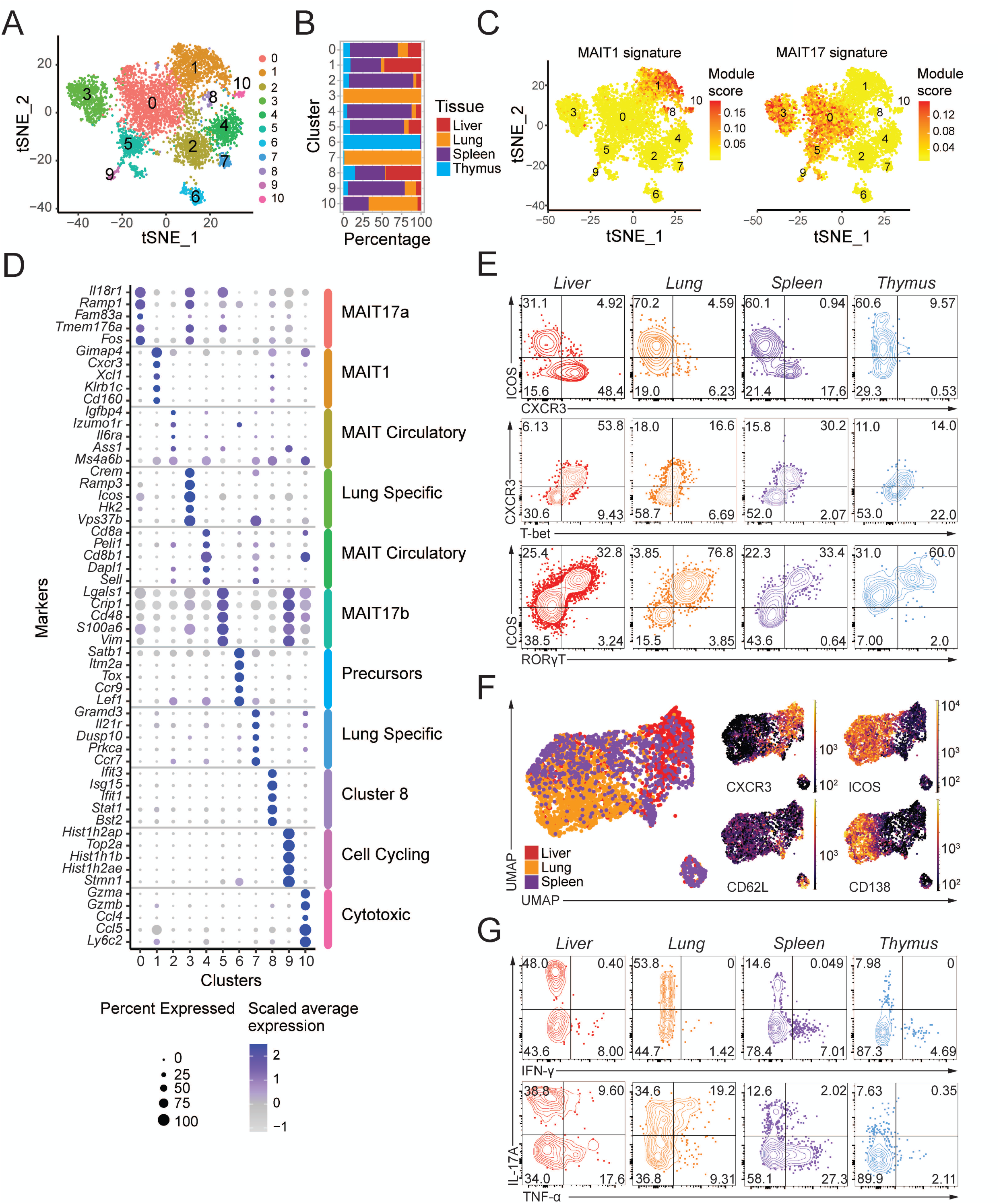
Heterogeneity of mouse MAIT cells. (A) Transcriptomic analysis of 6,080 mouse MAIT cells at steady state was performed using 10X Genomics platform. t-distributed stochastic neighbor embedding (t-SNE) plots were generated by pooling four individual scRNA-seq libraries of MAIT cells from thymus, lung, liver and spleen. Clusters were identified by shared nearest neighbor modularity optimization-based clustering algorithm. (B) Bar graph shows for each cluster the tissue origin of the MAIT cells contributing to that cluster. (C) Dot plot showing top 5 positive marker genes in each cluster. Color gradient and dot size indicate gene expression intensity and the relative proportion of cells within the cluster expressing each gene, respectively. (D) t-SNE plot showing the MAIT1 and MAIT17 signature scores for each cell. Signature scores are the difference between the average expression levels of a gene set and control genes for each cell. (E) Representative flow cytometry plots showing surface expression of CXCR3 and ICOS on MAIT cells from the indicated organs (top row) and the co-expression of T-bet with CXCR3 (middle) and RORγT with ICOS (bottom). (F) Flow cytometry data were acquired using a panel of 17 fluorescent parameters. MAIT cell data from liver, lung and spleen were used to perform UMAP dimensional reduction and unsupervised clustering using the FlowSOM algorithm on the OMIQ software. A total of 3791 MAIT cells were included in this analysis. (G) Cytokine expression by MAIT cells upon PMA/Ionomycin stimulation in vitro. Intracellular cytokine staining data are representative of 3-4 mice per group, representative of 2-3 experiments.

MAIT cell transcriptomes in different clusters could be associated with function as well as preferential tissue location. We used a gene signature for MAIT1 and MAIT17 phenotypes, based on an earlier RNA microarray of MAIT cells^21^, to define clusters enriched in these signatures. This analysis showed that MAIT cells in clusters 0, 3, 5 and 9 were enriched for MAIT17 signature (**Fig. 1C**). For example, the most differentially expressed genes (DEG) in MAIT cells of the largest cluster (cluster 0), including *Il18r1, Ramp1, Fos and Tmem176A* **(Fig 1D and Supplementary Table 1),** also were highly expressed in **IL-17 producing iNKT cells (NKT17 cells)**^32, 20, 33, 34^. On the other hand, Cluster 1 MAIT cells were enriched in Th1/NKT1 genes^32, 20, 33, 34^ including *Cd160*, *Klrb1c, Xcl1, Cxcr3 and Gimap4* **(Fig. 1D).** Additional MAIT cell clusters were distinguished by the expression of cell cycle genes or genes related to cytotoxicity (**Fig. 1D**). Cluster 2 and 4 were not enriched for either MAIT1 or MAIT17 but express circulatory cell markers such as *Sell*.

We used high-parameter flow cytometry and dimensional reduction to validate the phenotypic heterogeneity of MAIT cells. The majority of mouse MAIT cells could be divided into two populations based on the expression of markers defined from our transcriptomic analysis, ICOS and CXCR3. While ICOS expression strongly correlated with the expression of RORγT, suggesting ICOS marks MAIT17 cells **(Fig. 1E, 1F and Extended Data Fig. 1C)**, CXCR3 correlated with T-bet, suggestive of a MAIT1 population **(Fig. 1E, Extended Data Fig. 1C)**. As expected from the transcriptomic analysis, flow cytometry indicated that the ICOS^+^ MAIT17 cells were represented in all four organs **(Fig. 1F and Extended Data Fig. 1D)**. Of note, MAIT17 cells (CXCR3^-^, CD62L^-^) could be further divided based on Syndecan-1 (SDC1/CD138) expression into MAIT17a cells (SDC1^+^), and MAIT17b (SDC1*^-^*) populations (**Fig 1F, Extended Data Fig. 1E**). Selective Syndecan-1 has been reported to negatively regulate other innate-like IL-17-producing T cells, including iNKT cells and γδ T cells^35^ but in binding to extra cellular matrix proteins, Syndecan-1 expression also could be important for tissue maintenance. Both MAIT17 subsets had the highest prevalence in lung tissue, while CXCR3^+^ MAIT1-like cells were present predominantly in liver **(Fig 1E, 1F, Extended Data Fig. 1D, 1F)**. This distribution is similar to the iNKT cell functional subsets in different sites^32^.

We tested the functional capacity of these MAIT cell subsets by measuring cytokine production following stimulation. There were differences in the degree of activation of cells from different tissues, but MAIT cells capable of producing IL-17 were found in all organs, including the thymus (**Fig. 1G**). IFNγ and TNF producing cells were most prevalent in the spleen and liver, consistent with the scRNA-seq and flow cytometry indicating enrichment for MAIT1 cells in these sites, but some MAIT thymocytes also produced these cytokines. Together, these data indicate a strong correlation between transcriptomic, phenotypic, and functional data in defining MAIT1, heterogenous MAIT17 states and some subsets that did not fit into either category. The different subsets were present to varying degrees in all tested tissues, with the exception of the lung-specific and thymic precursor subsets.

### Large-scale shifts in gene expression by MAIT thymocytes

The data demonstrated that MAIT cells from mouse thymus are found in most of the clusters with peripheral MAIT cells **(Fig. 1B and Extended Figure 1B)**. These results, along with the activation assay results (**Fig. 1G**), confirmed the previously reported presence of mature, functional MAIT cells in the thymus^9, 10, 16, 21^. It has been reported that immature thymic CD24^+^CD44^−^ stage 1 precursor cells transition to stage 2 (CD24-CD44-) and give rise to mature CD24^−^CD44^+^ MAIT1 and MAIT17 cells^9^. To create an unbiased model of MAIT cell thymic differentiation that encompasses the different clusters, we have employed the Monocle 2/DDRtree algorithm^36^ to enable pseudo-time ordering based solely on the scRNA-seq of the MAIT cell differentiation stages of thymus cells **(Fig. 2A, B)**. The analysis indicates that relatively immature or precursor thymic MAIT cells divide into two branches: one leading to MAIT1 thymocytes and another leading to MAIT17 thymus cells **(Fig. 2B)**. Hierarchical clustering of gene expression of the thymic MAIT cell clusters generated 12 gene modules of genes that tended to be co-expressed in individual cells along the trajectory (**Fig. 2C and Supplementary Table 2).**

**Figure 2:**
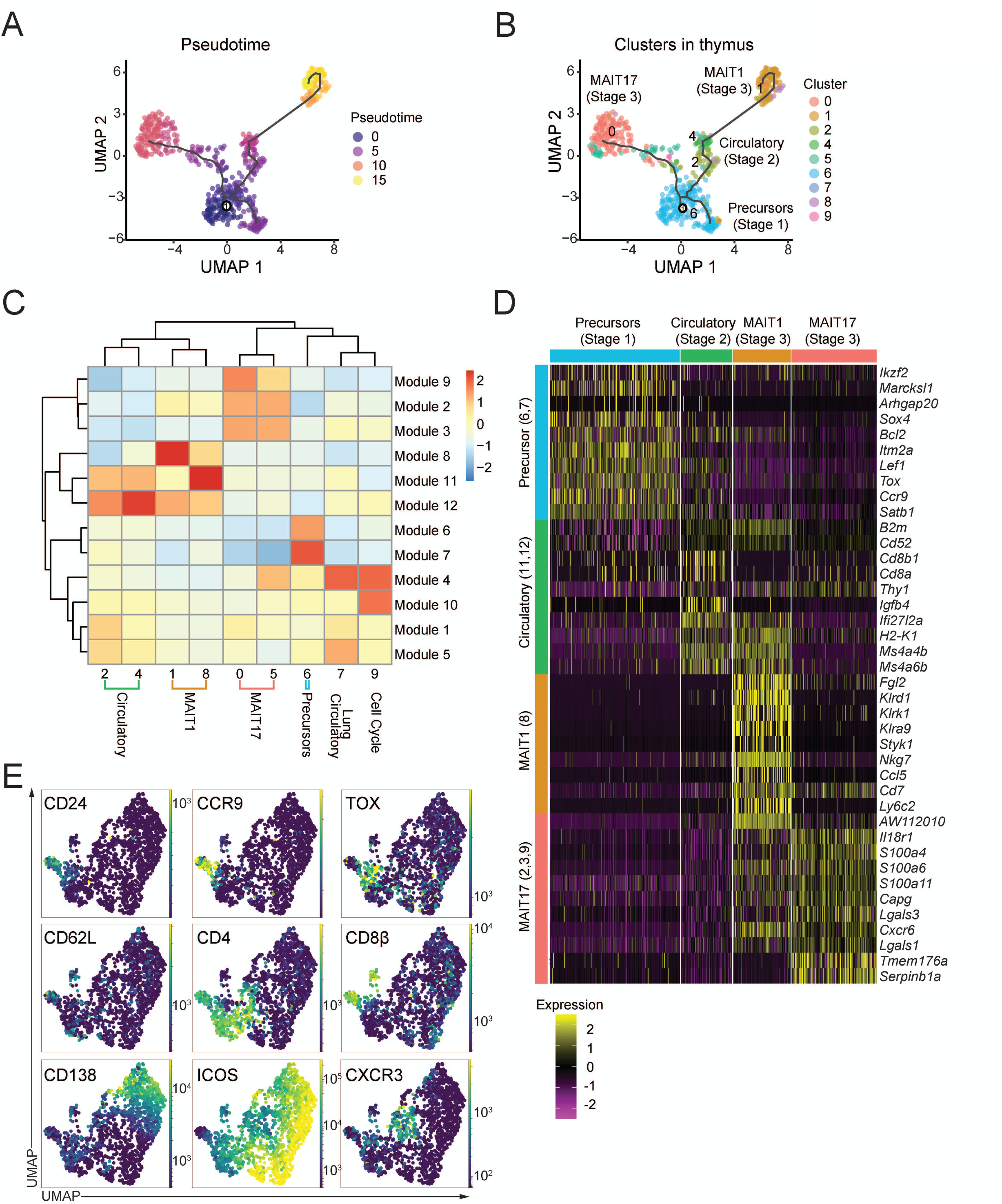
MAIT cell changes in gene expression during thymus differentiation. (A) Single-cell trajectories of mouse MAIT thymocytes constructed using Monocle 3. UMAP shows cells colored by pseudotime values along the trajectory. (B) UMAP showing distribution of thymic MAIT cell clusters across branches of single-cell trajectories. Cluster colors and numbers as in Fig 1A. (C) Heatmap showing different stages of thymus MAIT cell differentiation and respective cell clusters on the *x*-axis and co-regulated gene modules on the *y*-axis. Modules consist of genes that are differentially expressed along the thymus trajectory path. The legend shows color-coded aggregate scores for each gene module in all the clusters; positive scores indicate higher gene expression. (D) Scaled average expression heatmap of top 10 genes from each thymocyte gene module based on high Morans I value that were expressed in the indicated clusters of MAIT thymocytes. (E) Flow cytometry data were acquired using a panel of 17 different fluorescent parameters. MAIT cell cytometry data from mouse thymi (n=5) were used to perform UMAP dimensional reduction and unsupervised clustering using the FlowSOM algorithm on the OMIQ software. A total of 1,568 MAIT thymocytes from 9-week-old mice were used for the analysis. All mice were 9-week-old C57BL/6 females.

Based on the expression of key genes, we could align the gene expression modules with the stages of MAIT cell maturation derived earlier from flow cytometry and functional assays. MAIT cells in cluster 6 have high expression of genes in two modules (6 and 7, precursor modules), which include *Satb1, Ccr9, Tox, Lef1, Itm2a, Bcl2* and *Sox4* **(Fig. 2D, Extended Data Fig. 2B, Extended Data Fig. 2C, Supplementary Table 2)**. Therefore, this cluster served as the starting point for the pseudotime analysis. Flow cytometry analyses confirmed co-expression of TOX protein by cells that co-express typical stage 1 genes, such as CD24 and CCR9 **(Fig. 2E)**, along with the absence of CD44 (data not shown). In addition, CD44^-^ cells have the highest expression of SATB1 and LEF1 on the protein level **(Extended Data Fig. 2D**), further validating its identity as a group that contains relatively immature thymic MAIT cells or stage 1 cells. Cells from clusters 2 and 4 lack expression of both *Cd24a*, typical of the most immature cells and *Cd44,* which marks mature or stage 3 cells **(Extended Data Fig. 2A)** while the pseudotime trajectory indicates they are closer to the precursor population. Furthermore, several genes from the precursor modules, such as *Lef1* and *Satb1 (Id3 not shown)*, are expressed to some extent by these putative stage 2 MAIT cells (**Fig. 2D, Supplementary Table 2).** Altogether, this suggests that these cell clusters contain intermediate or stage 2 MAIT cells. These stage 2 MAIT cell clusters share two gene modules (11 and 12, Stage 2 modules), that include expression of *Ms4a4b, Ms4a6b and Ccr7* **(Fig. 2D, Extended Data Fig. 2B and Extended Data Fig. 2C, Supplementary Table 2)**. These clusters could be distinguished by the expression of *Cd4* by cluster 2 and *Cd8a*, *Cd8b1*, and *Klrd1* by cluster 4. CD4 expression was reported to be enriched in stage 1 and stage 2 thymic MAIT cells, while CD8 is increased in stage 3 MAIT thymocytes^8, 37^.These findings and the single-cell trajectory analysis suggest that of the two stage 2-like MAIT cell clusters, cluster 4 is more mature and perhaps closer to MAIT1 cells. Separate groups of CD4 and CD8α stage 2 MAIT thymocytes were confirmed by flow cytometry **(Fig 2E).**

Stage 3 MAIT1 thymocytes have high expression of gene module 8 including several natural killer cell receptors, such as *Klrd1 and Nkg7*, whereas in MAIT17 cells, expression of genes modules 2, 3 and 9 were increased and included transcripts such as *Il18r1, Lgals3 and Tmeme176a* **(Fig. 2D, Extended Data Fig. 2B, Extended Data Fig. 2C, Supplementary Table 2)**. A recent study^38^ provided data indicating that NKT17 cells are generated earlier in the thymus than NKT1 cells, which also seems to be true for our *in-silico* analysis of thymic MAIT17 and MAIT1 cells, with MAIT17 cells closer to the precursor stages (**Fig. 2B**). Taken together, these data show that the stages defined on the basis of the expression of CD24 and CD44 in fact encompass the MAIT cells in the thymus. When an unbiased analysis is undertaken, that the stage 3 cells are almost entirely MAIT1 and MAIT 17 cells in C57BL/6 mice, and the stage 2 classification is heterogenous and possibly containing cells with different potentials and/or different degrees of differentiation.

### MAIT circulatory and precursor cells

MAIT cells have been reported to be tissue-resident cells in mice^20^. A tissue resident gene signature was most enriched in the lung cell cluster 3 (**Fig. 3A**). MAIT cells can be found in the circulation, and they are abundant in human peripheral blood^8^. Using a gene expression signature score for circulatory mouse CD8^+^ T lymphocytes^39^, we found that peripheral MAIT cells similar to thymus stage 2 cells (clusters 2, 4) and a smaller population of lung-specific cells (cluster 7) were distinguished by enrichment of circulatory signature genes **(Fig. 3A)**. MAIT cells in these clusters expressed *Ccr7, Sell* (encoding CD62L) and *Lef1* **(Fig. 3B)** and were most prevalent in the spleen, but detectable in different tissues. (**Fig. 1B**). Cluster 7 circulatory-like MAIT cells also exhibited expression of some tissue-resident genes, unlike clusters 2 and 4. Importantly, cells in all three of these putative circulatory MAIT cell clusters, even those in the periphery, had relatively low expression of mRNA encoding *Cd44*, *Rorc* and *Tbx21* **(Fig, 3B)**, suggesting they could be less mature and more similar to stage 2 MAIT thymocytes. We refer to these relatively immature cells with enriched circulatory gene expression signature as circulatory/precursors (MAIT_CP_). We applied Monocle analysis to the scRNA-seq data to perform pseudotime trajectory analysis of spleen and lung MAIT cells. Consistent with our designation, in spleen, cluster 2 is at the root of the trajectory **(Fig. 3C and Supplementary Table 3)** whereas in lung the root of the trajectory consists mainly of cells from cluster 7 **(Fig. 3D and Supplementary Table 4)**.

**Figure 3:**
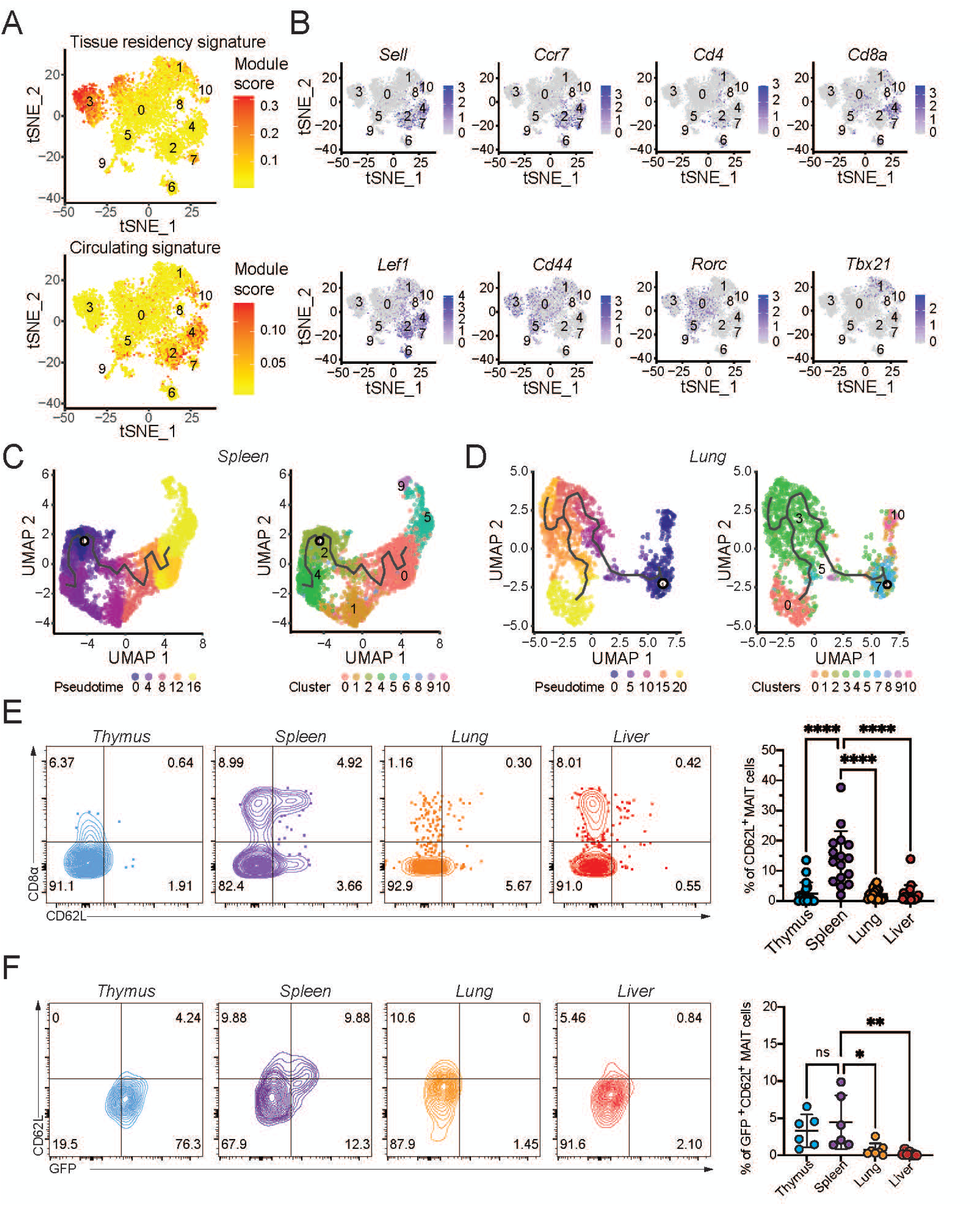
Circulatory MAIT cells are related to recent thymic emigrants. (A) t-SNE showing the circulatory and tissue resident signature scores for each cell MAIT cell from the four organs from the mouse scRNA-seq data. (B) t-SNE showing expression of *Sell, Ccr7, CD4, CD8a, Lef1, CD44, Rorc* and *Tbx21* transcripts in MAIT cells. Single-cell trajectory analysis of MAIT cells in spleen (C) and lung (D) showing cells ordered in pseudotime and placed along a trajectory of gene expression changes, constructed using Monocle 3. Figure shows the UMAP with cells colored by pseudotime values (left) and the UMAP with cells colored by clusters as in Fig. 1A (right). Darker violet color denotes the root cells and yellow color denotes the outcome. (E) Representative flow cytometry (left) plots showing surface expression of CD62L and CD8α by MAIT cells in C57BL/6 mice in the indicates tissues. Percentage of CD62L^+^ MAIT cells (right, displayed as mean□□SD) in the indicated organs (n = number of mice). One-way analysis of variance (ANOVA) with post-hoc Tukey test. Liver n= 19, Lung n= 16, Spleen n = 16, Thymus n= 19. ****: adjusted p-value < 0.0001. Mixed male and female mice, 14.8±6.2 weeks-old. (F) Representative flow cytometry plots (left) showing surface expression of CD62L and GFP by MAIT cells in four-week-old Rag2:GFP mice (n=6) in the indicated tissues. Percentage of GFP^+^CD62L^+^ MAIT cells (right, displayed as mean SD) in the indicated organs. One-way ANOVA with post-hoc Tukey test, *P□<0.05, **P□<0.01. Flow cytometry data are representative of 2-3 experiments.

Cells in clusters 2, 4 and 7 also express increased amounts of genes encoding ribosomal proteins **(Extended data Fig. 3A)**, suggestive of proliferative capacity. By flow cytometry, we confirmed that MAIT cells expressing CD62L were most abundant in the spleen, especially in CD8^+^ MAIT cells (**Fig. 3E**) and they are also prevalent in the blood **(Extended Data Fig. 3B)**, in accord with the hypothesis that they circulate through the vasculature. The spleen trajectory indicates that the cluster 4 cells are slightly more mature than those in cluster 2, which fits with the thymus trajectory analysis **(Fig. 2B**), while MAIT17b cells lacking Syndecan-1 were more mature than MAIT17a (cluster 0) (Fig.3C).

The presence of perhaps relatively immature MAIT_CP_ cells in the periphery led us to identify MAIT cells that might be recent thymic emigrants (RTEs). We used transgenic mice that express green fluorescent protein (GFP) under the control of the recombination-activating gene 2 (*Rag2*) promoter (Rag2:GFP). In these mice, GFP expression indicates cells that recently rearranged their TCR genes^40, 41^ providing an indicator of the timing of initiation of TCR expression and maturation. In the Rag2 reporter mice, the pattern of CD62L expression by MAIT cells was not altered, with more expression in spleen (**Fig. 3F**) and blood **(Extended Data Fig. 3C)**, a pattern similar to WT controls. The data showed that a high percentage of MAIT thymocytes in four-week-old mice expressed the Rag2 reporter **(Fig. 3F**), suggesting that MAIT cells in young mice have recently rearranged *Tcra*. The spleen contained a significant number of MAIT cells with co-expression of CD62L and Rag2:GFP, indicating the presence of a relatively immature, peripheral population, while lung and liver had a lower number of these cells.

An earlier report showed that CCR7^+^ iNKT cell precursors^42^, which also expressed LEF1, egress from the thymus and undergo final maturation in the periphery. Some of the thymic MAIT cells also expressed CCR7, and they are mostly RAG2:GFP^+^. The spleen also contained a significant proportion of GFP^+^ MAIT cells, which were predominantly CCR7^+^ (**Extended Data Fig. 3D)** consistent with the hypothesis that MAIT cell RTEs are most prevalent there. Expression of LEF1 was higher on CD62L^+^ thymic MAIT cells as compared to their more mature, CD44^+^ MAIT cell counterparts **(Extended Data Fig. 3E)**. Furthermore, the thymus had the highest percentage of LEF1^+^ MAIT cells as compared to peripheral MAIT cells **(Extended data Fig. 3F)**. Overall, expression of the RAG2 reporter, CCR7, CD62L were correlated in the thymus and spleen. The data therefore are consistent with a model in which cell types similar to stage 2 MAIT thymocytes are in RTE that circulate in the blood and are in the spleen. We therefore propose that MAIT_CP_ are circulatory MAIT cells that are precursors for cells that further differentiate in the periphery.

### Mouse MAIT17 cells are metabolically active

Following activation and differentiation, CD4^+^ and CD8^+^ T cells profoundly change their cellular metabolism. Effector-like as opposed to memory-like and tissue-resident states are controlled by divergent metabolic programs, relying on glycolytic versus mitochondrial or fatty acid oxidation metabolism, respectively^31^. Analysis of the scRNA-seq signature scores for oxidative phosphorylation, mitochondrial genes, fatty acid metabolism and glycolysis showed that MAIT17 clusters had the most enrichment for oxidative phosphorylation genes, with the least expression in MAIT_CP_. **(Extended Data Fig. 4A)**. To further analyze the metabolism of heterogenous MAIT cells based on scRNA-seq, we used the Compass algorithm^43^, which computes the analysis of variance (ANOVA) for each reaction in different metabolic pathways. We selected the reactions labeled by the Recon2 database^44^ as involved in glycolysis/gluconeogenesis, citric acid cycle, fatty acid oxidation or amino acid metabolism pathways. When compared to the thymocyte precursors (cluster 6), MAIT17 clusters showed the highest Cohen’s D score for each pathway, followed by MAIT1, with lower Cohen’s D scores in the MAIT cell lung tissue-resident cluster 3 and MAIT_CP_ **(Extended Data Fig. 4B and Supplementary Table 5)**. This analysis is therefore consistent with the gene signatures in showing a higher metabolic pathway expression in MAIT17 cells versus reduced metabolism in MAIT1 and MAIT_CP_.

Because transcriptomic data do not always accurately reflect metabolic activity, we measured MAIT cell metabolic parameters by flow cytometry to validate these findings. We quantified the uptake of fatty acids and glucose, the cytoplasmic lipid droplet content, and the activity of mitochondria with membrane-potential sensitive MitoTracker Deep Red FM, which accumulates in active mitochondria. Large differences were evident during the differentiation of total MAIT cells in thymus, with stages 1 and 2 cells, gated as in **Extended Data Fig. 5a**, significantly less active metabolically compared to stage 3 for all measures. The predominant mature MAIT cells in the thymus are MAIT17 cells, and compared to MAIT1 cells, MAIT17 thymocytes showed higher levels for all the metabolic parameters, except glucose uptake **(Fig 4A and 4B)**. Consistent with metabolic reaction modeling, MAIT_CP_ were significantly less active than MAIT17 and in that regard comparable to MAIT1 cells. These data indicate that thymic MAIT17 cells adopt a distinctly active metabolic phenotype during functional differentiation.

**Figure 4:**
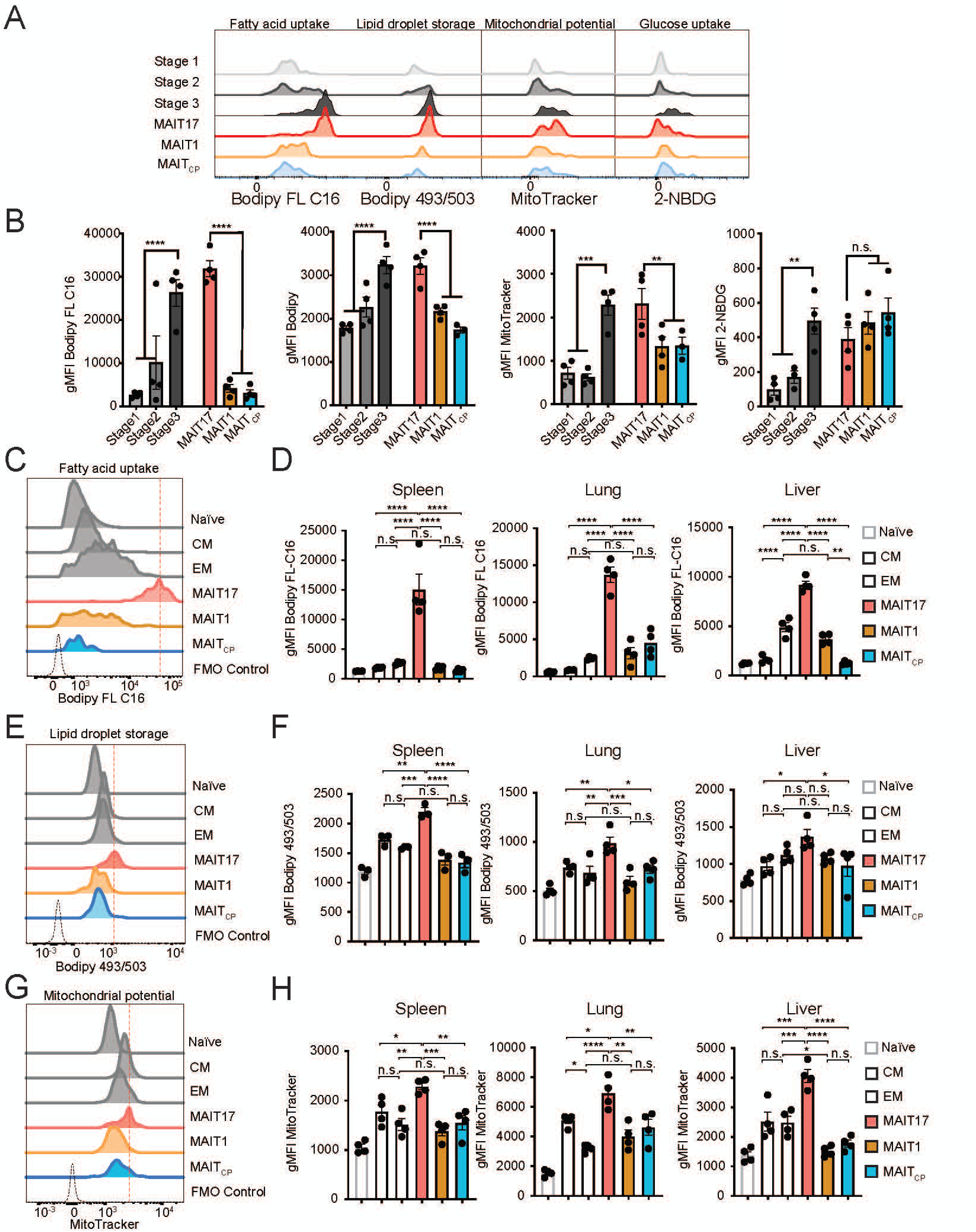
Mouse MAIT cell subsets have distinct metabolic features. (A-B) Metabolic parameters of MAIT thymocytes were quantified for the MAIT cell differentiation stages 1-3 and MAIT1, MAIT17 and MAIT_CIRC_ subsets. Representative histograms (A) and quantification (B) are depicted as geometric mean fluorescence intensity (gMFI). Thymic MAIT cell precursor stages 1-3 were defined based on CD24 and CD44 expression. Neutral lipid droplets were quantified by Bodipy 493/503 fluorescence (left), fatty acid uptake was quantified as intensity of Bodipy FL C16 fluorescence (center left), mitochondrial content was quantified as Mitotracker Deep Red FM fluorescence (center right) and glucose consumption by uptake of 2-deoxy-2-[(7-nitro-2,1,3-benzoxadiazol-4-yl)amino]-D-glucose (2NBDG) (right). (C-H) Cells were isolated from spleen, lung and liver and metabolic parameters were quantified in CD8^+^ T cells and MAIT cell subsets. TCRβ^+^ CD8^+^ T cells excluding MAITs were subdivided into naïve, central memory (CM) and effector memory (EM) subsets based on expression of CD62L and CD44. Quantification (C) and representative histograms (D) of fatty acid uptake in the indicated cell types and organs were measured as gMFI of Bodipy FL C16. Quantification (E) and representative histograms (F) of neutral lipid droplet content in indicated cell types and organs were measured as gMFI of Bodipy 493/503 fluorescence. Quantification (G) and representative histograms (H) of mitochondrial content in indicated cell types and organs were measured as gMFI of MitoTracker Deep Red FM signal. Data from 3-4 mice per group, representative of ≥3 experiments. Data analyzed by one-way ANOVA with Dunnett’s post-test for multiple comparisons, displayed as mean±*□□*SEM, \**P□*<0.05, \*\**P□*<0.01 \*\*\**P□*<0.001 and \*\*\*\**P□*<0.0001.

We compared peripheral mouse MAIT cells to CD8^+^ naïve, central memory (CM) and effector memory (EM) T cells, gated as in **Extended Data Fig 5A**. As in the thymus, there was heterogeneity comparing subsets: MAIT17 cells in all sites had significantly elevated uptake of fatty acids, lipid content, and mitochondrial potential compared to MAIT1, MAIT_CP_ or the CD8^+^ memory T cell subsets **(Figure 4C-H)**. In contrast, a time course analysis showed that MAIT1 cells have higher glucose uptake compared to MAIT17 cells in liver and spleen, although this was not reflected in the scRNA-seq data **(Extended Data Fig. 5B)**. Regardless, these data suggest that all MAIT cells engage in fatty acid uptake, fat storage and mitochondrial membrane polarization, suggesting they may depend on mitochondrial respiration to generate ATP. However, our data suggest that MAIT1 cells support the generation of ATP through consumption of glucose, while MAIT17 cells preferentially metabolize fatty acids. Importantly, peripheral MAIT_CP_ closely mirrored the metabolic program of their CD62L^+^ thymic counterparts. Together, this suggests that adoption of differential metabolic program by MAIT17 cells occurs in thymic MAIT subsets and depends on the functional differentiation they undergo in the thymus or periphery rather than their ultimate tissue localization.

### Heterogeneity of human MAIT cells

In order to determine the extent of human MAIT cell heterogeneity and to assess the homology of human and mouse MAIT cell subsets, we carried out scRNA-seq of sorted human MAIT cells from thymus, peripheral blood and lung. Thymus tissues were obtained from children undergoing partial thymectomy due to cardiac surgery. Lung and peripheral blood were obtained from the same adult donors, undergoing surgery for early-stage lung cancer^45^. MAIT cells were identified as Vα7.2^+^, human MR1 5-OP-RU tetramer^+^ cells as shown in **Extended Data Fig. 6A**. Human MAIT cells were also heterogenous **(Fig. 5A and Fig. 5B and Supplementary Table 6)** and most MAIT cell clusters were highly tissue-specific **(Extended Data Fig. 6B).** Importantly, demultiplexing analysis indicated that the clusters contained cells from multiple donors **(Extended Data Fig. 6C)**. There were multiple thymus-specific clusters **(Fig. 5B and Extended Data Fig. 6B)** and one nearly completely lung specific subset (cluster 4). Cluster 0 consisted mostly of cells from PBMCs and these cells expressed MAIT1 cell genes such as *NCR3, KLRB1 and GZMK* **(Fig. 5C)**. Like PMBCs, most clusters from lung and cluster 2 from thymus showed enrichment of MAIT1 signature score. In addition to cluster 4, clusters 3 and 5 were also enriched for lung MAIT cells, and these clusters were enriched for MAIT17 signature score, although this did not exclude co-expression of MAIT1 genes **(Fig. 5D)**. Similar to the mouse, however, MAIT cells in these lung MAIT cell clusters were enriched for a tissue-residency signature gene, whereas a circulatory gene expression signature was detected in MAIT cells from PBMCs **(Fig. 5D)**. Previous work showed lung lymphocytes from mice also had increased expression of genes associated with activation, such as members of the *NFKB* and *AP1* families^46^. Human lung MAIT cells also showed increased expression of genes associated with activation, including *TNFAIP3* and *FOS* **(Fig. 5C)**.

**Figure 5:**
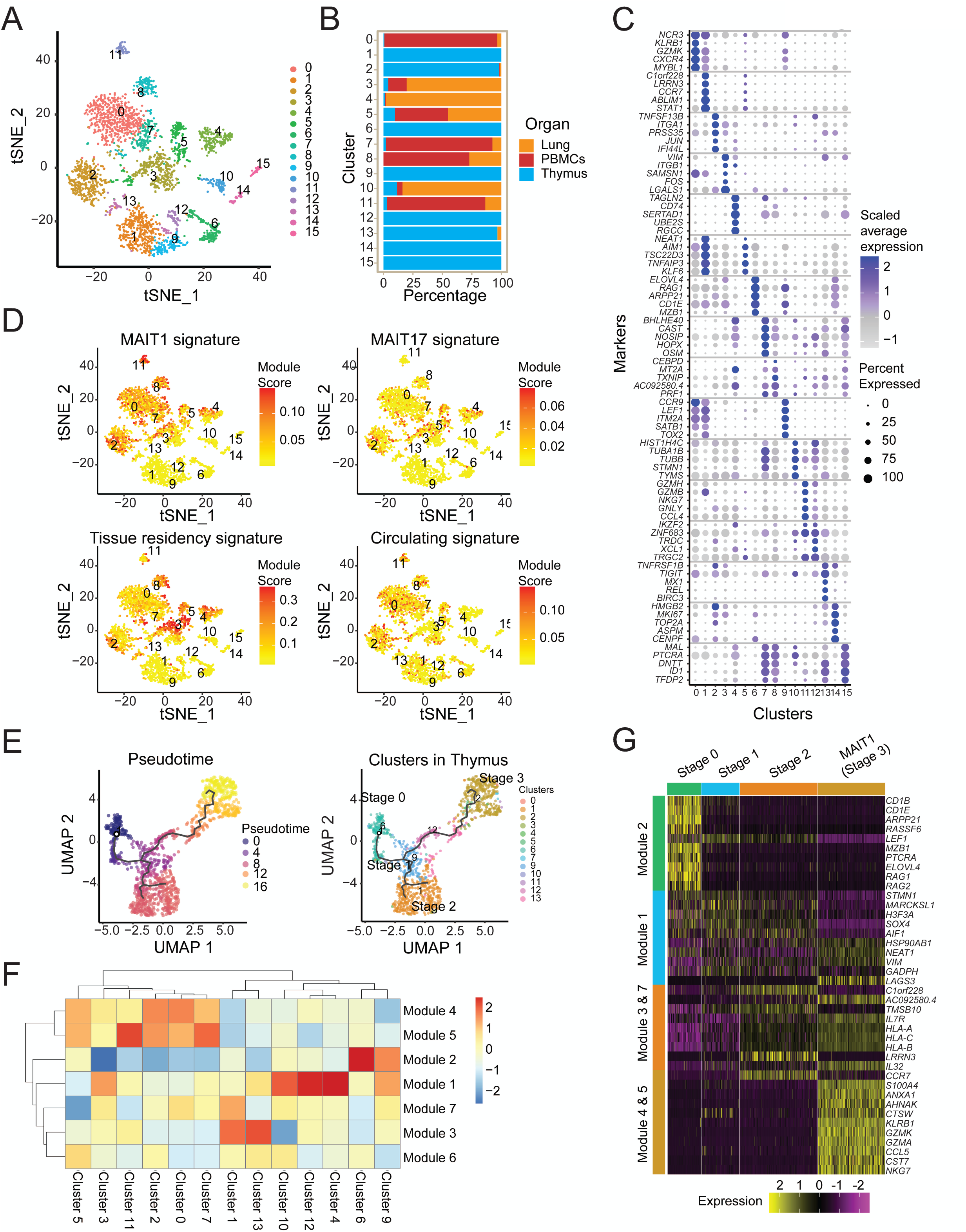
Heterogeneity of human MAIT cells. (A) Transcriptomic analysis on 3,020 human MAIT cells (hMR1:5-OP-RU^+^ TCRβ^+^) was performed using the 10X Genomics platform. t-SNE plots generated by combining three individual scRNA-seq libraries from human thymus (n=5), lung (n=4) and PBMCs (n=4). (B) Bar graph shows the contribution of MAIT cells from different tissues to individual clusters. (C) Dot plot showing top 5 positive marker genes in each cluster. Color gradient and size of dots indicate gene expression intensity and the relative proportion of cells within the cluster expressing each gene, respectively. (D) t-SNE showing the MAIT1, MAIT17, tissue residency and circulating signature scores for each cell. Positive scores indicate high expression of genes in the gene set of interest as compared to randomly selected controls. (E) UMAP (left) of human MAIT cells from thymus with cells ordered in pseudotime and UMAP showing distribution of thymic MAIT cells (right) across branches of single-cell trajectories. Cells are colored and numbered by clusters as in Fig. 5A. (F) Heatmap showing different stages of development and respective cell clusters on the *x*-axis and co-regulated gene modules on the *y*-axis. Modules were generated with genes that are differentially expressed along the trajectory path. The legend shows color-coded aggregate module scores for gene modules for cells in each cluster; positive scores indicate higher gene expression. (G) Scaled average expression heatmap of top 10 genes from modules that were expressed in indicated stages of MAIT cell development based on high Morans_I value as shown in Fig 5F. These genes were selected based on their expression changes as the cells progress along the MAIT cell developmental trajectory.

Clusters from the thymus provide evidence for differentiation from immature cells leading to a mature MAIT1 cell population (cluster 2). The most immature cluster, cluster 6 or here called MAIT0 cells, expressed genes such as *RAG1, RAG2, PTCRA, LEF1, CD1E* and *CD1B* **(Fig. 5C)**. This agrees with an earlier report using bulk RNA-sequencing of human MAIT thymocytes^10^, MAIT cells in this cluster appear to be even less mature than typical mouse thymus stage 1 cells. Cluster 9 MAIT cells expressed *CCR9, LEF1, ITM2A, SATB1* and *TOX* **(Fig. 5C)** while lacking the expression of *CD27* and *KLRB1* (encoding CD161) **(Extended Data Figure 6D)**, and therefore are similar to the previously defined stage 1 cells. To investigate the gene expression dynamics underlying the human MAIT cell differentiation program, we created a pseudo-time ordering for thymus MAIT cell transcriptomes **(Fig. 5E)**. This analysis allowed definition of modules of genes that were co-expressed for each cluster **(Fig. 5F and Supplementary Table 7)**. The analysis revealed a precursor gene module (module 2) found in immature cell clusters 6 and 9 **(Fig. 5G and Extended Data Figure 7A).** There are also MAIT1 gene modules (4 and 5), characterized by increased expression of genes such as *NKG7* and genes encoding granzymes **(Fig. 5G, Extended Data Figure 7A and 7B)**. Pseudotime analysis suggests that cluster 6 containing the Stage 0 cells gives rise to cluster 9, an intermediate transcriptional state (stage 1), which further differentiates into a stage 2-like cells (cluster 12) with *CD27* expression. This cluster then branches into stage 3-like cells (cluster 2) that express *KLRB1*. A separate branch can give rise to cluster 1, enriched in gene modules 3 and 7. This MAIT cell cluster has higher expression of circulatory or precursor transcripts, such as *LEF1* and *CCR7* **(Fig. 5G, Extended Data Figure 7A and 7B),** and also *LRRN3*, which is co-expressed with *LEF1* and *CCR7* and marks naïve human T cells^47^. These data are consistent with the notion that this group of human MAIT cell clusters are not fully differentiated and similar to mouse stage 2 thymocytes and MAIT_CP_, although their differentiation potential is unknown.

### Human MAIT cells have increased fatty acid uptake and storage

We calculated metabolic signature scores for the transcriptomes of human MAIT cell clusters **(Extended Data Fig. 8A).** We also functionally tested the metabolic activity of human circulatory (blood) and tissue (lung) MAIT cells and compared them to naïve, effector and memory CD8^+^ T cell populations (gated as in **Extended Data Fig. 8B)**. As expected, our data indicate that the CD8^+^ memory T cell subsets were higher for their metabolic readouts compared to naïve or effector T lymphocytes **(Fig. 6A-6D)**, which agrees with previous research^48^. As reported previously^49^, the CD8^+^CD103^+^ subset, a phenotype of tissue-resident memory T cells (TRM), had the most enhanced fatty acid uptake. We also analyzed MAIT cell phenotypic subsets, including the largest population of MAIT cells (CD103^-^, CD161^+^), and a smaller population of CD103^-^, CD161-MAIT cells. In human lung there also was a population of CD103^+^, CD161^+^ MAIT cells **(Extended Data Fig. 8B)**. The human MAIT cell subsets had a larger reservoir of stored lipids and also actively took up high amounts of fatty acids, comparable to TRM cells and even greater than central and effector memory cells **(Fig. 6A-6D)**. In contrast, the mitochondrial potential was low in all MAIT cell subsets, and was not significantly different from naïve or effector T cells. Therefore, the human MAIT cell metabolic parameters were not restricted to a phenotypic subset or to lung as opposed to PMBCs **(Fig. 6A-6D**), they resembled mouse MAIT17 cells in their high lipid stores and substantial uptake of fatty acids, did not have a higher mitochondrial potential, and therefore more similar in that regard to MAIT1 and MAIT_CP_.

**Figure 6:**
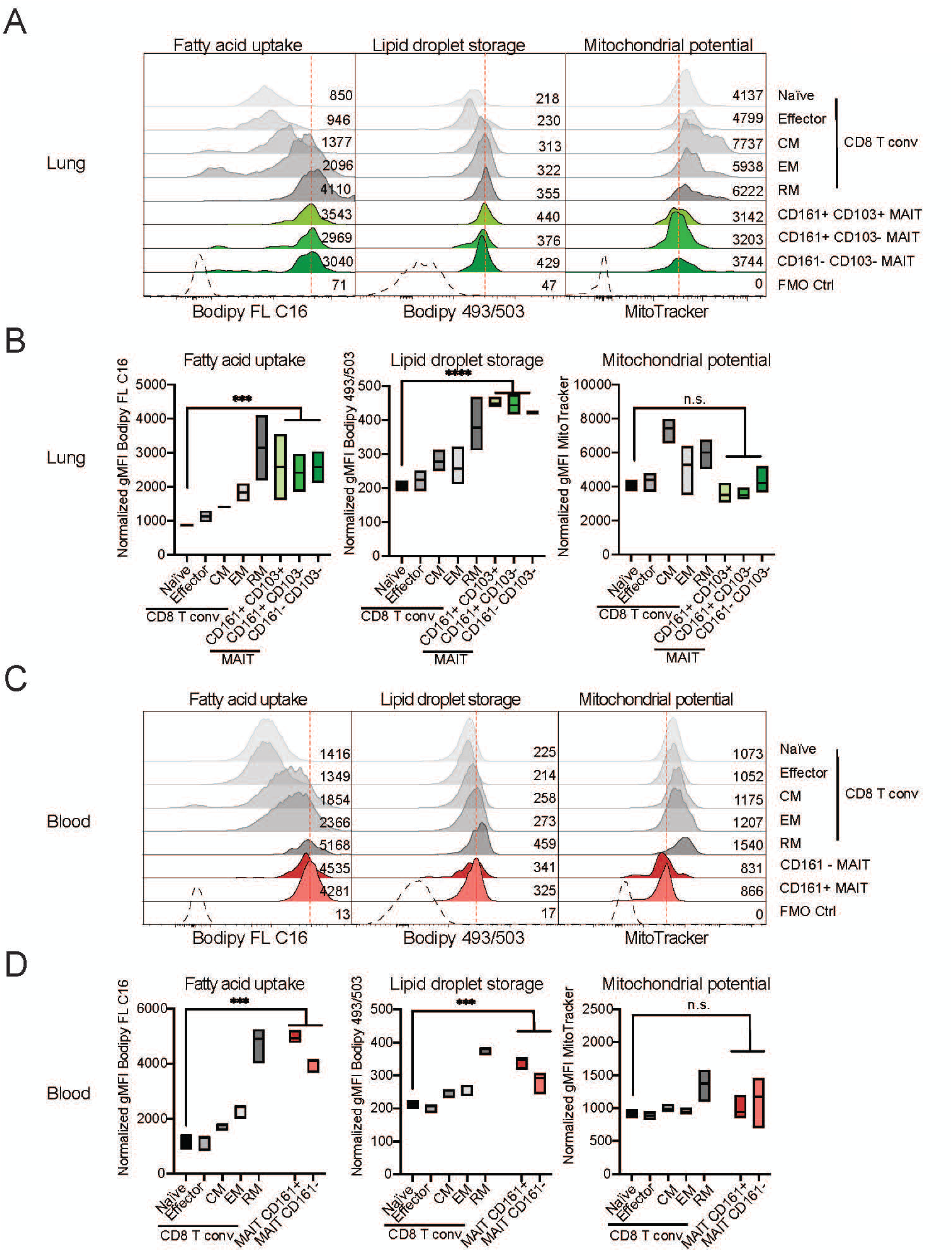
Human MAIT cell metabolic parameters differ from naïve CD8^+^ T cells. Cells were isolated from paired samples of human lung biopsies (A and B) or blood (C and D) and metabolic parameters were quantified in CD8^+^ T cell and MAIT cell subsets. TCRβ^+^ CD8^+^ T cells excluding MAITs were subdivided into naïve, central memory (CM), effector memory (EM) and resident memory (RM) subsets based on expression of CD45RA, CCR7 and CD103. Representative histograms (A, C) and quantification (B, D) of fatty acid uptake (left) was measured as gMFI of Bodipy FL C16. Neutral lipid droplet content (middle) was measured as gMFI of Bodipy 493/503 fluorescence. Mitochondrial potential is indicated as gMFI of Mitotracker Deep Red FM signal (right). Data combined from 2 experiments and 3 patients (A-B) or from 3 experiments and 3 donors (C-D). Data were analyzed by one-way ANOVA with Dunnett’s post-test for multiple comparisons, displayed as mean±□□SEM, *P□<0.05, **P□<0.01 ***P□<0.001 and ****P□<0.0001.

### Homology of human and mouse MAIT cell populations

To evaluate the similarities in the transcriptional signatures of human and mouse MAIT cell subsets, we performed integration of the human and mouse dataset^50^. Post-integration, 14 integrated MAIT cell clusters were identified, including some that were tissue specific **(Fig. 7A)**. Several clusters contained MAIT cells from both species **(Fig. 7B** and **Extended Data Fig. 8C**). This is particularly true for MAIT cell precursors in the thymus. MAIT cells in integrated or *i*-cluster 5 consisted of cells from precursors including mouse MAIT cluster 6 and human MAIT cluster 9 (**Extended Data Fig. 8C**) with expression of *ITM2A, CCR9 and TOX* and other genes characteristic of thymus differentiation **(Fig. 7C and Extended Data Fig. 9A)**. Cells in *i*-cluster 0 contained mouse and human MAIT thymocytes and also mouse spleen cells. These MAIT cells have a circulatory gene expression signature, including *SELL* and *IGFP4* **(Fig. 7C and Extended Data Fig. 9A)**, characteristic of MAIT_CP_ **(Fig 1D)**, indicating similarity between thymus stage 2 MAIT cell transcriptome is similar between the two species. Thymic MAIT cell subsets, however, did not completely overlap, as *i*-cluster 12 contained only stage 0 human MAIT thymocytes expressing *RAG1* and *RAG2* were not found in mice **(Fig. 7C and Extended Data Fig. 9A).**

**Figure 7:**
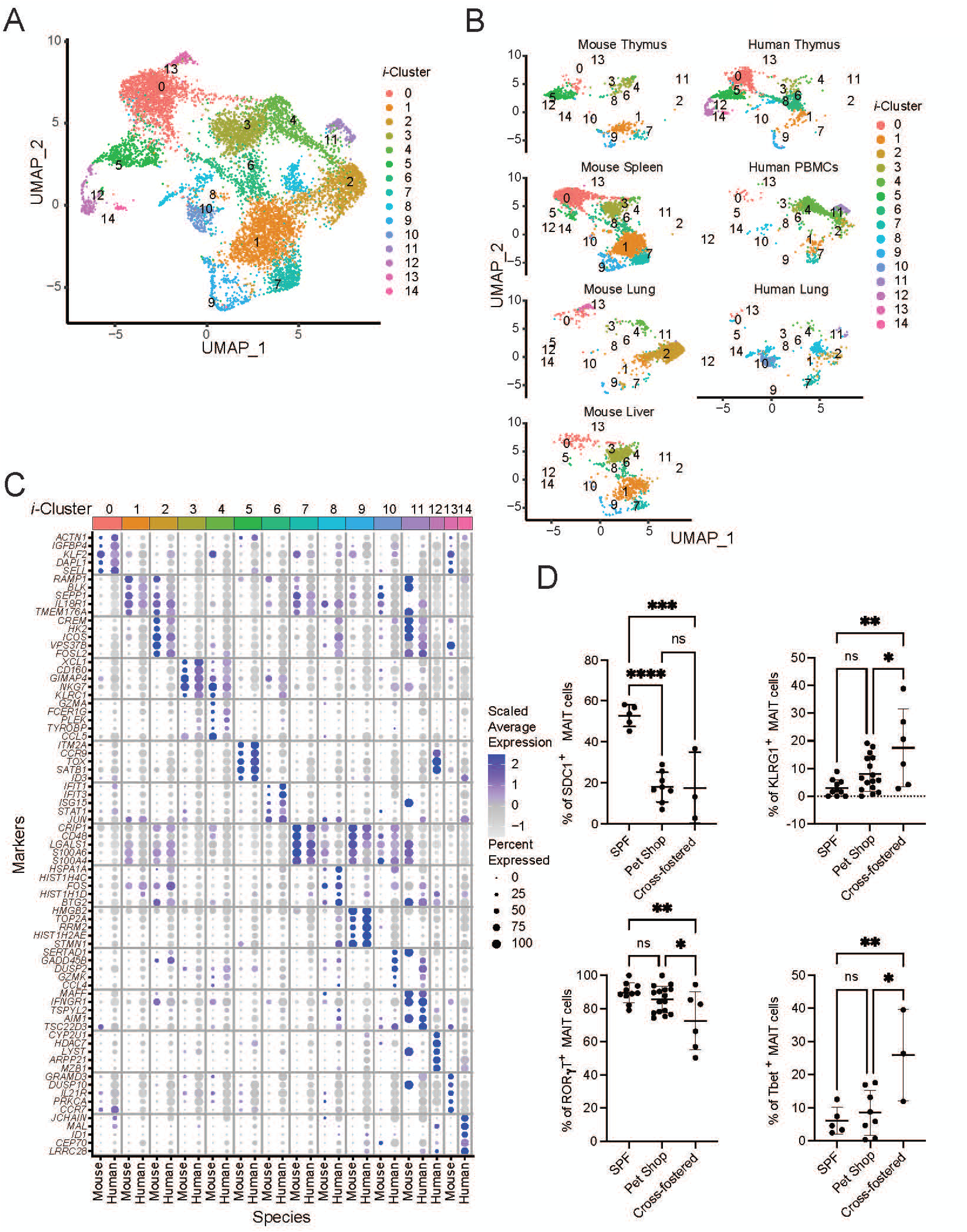
Divergent mouse and human peripheral MAIT cell subsets. (A) Aggregated UMAP representation of scRNA-seq data from mouse and human MAIT cells. (B) Mouse and human cells shown in separate UMAPs with the same coordinates as in Fig 7A. (C) Dot plot showing top 5 marker genes in each integrated cluster across both mouse and human cells. Color gradient and size of dots indicate gene expression intensity and the relative proportion of cells (within the cluster) expressing each gene respectively. (D) SPF mice were cross-fostered with mice from pet shops. The figure represents the percentage of MAIT cells expressing SDC1, KLRG1, T-bet and RORγT in lungs of the indicated mice. Data analyzed by one-way ANOVA with Tukey test displayed as mean± S.D. SPF mice n = 10, Pet shop mice n=16 and Cross-fostered mice n=6, *P□<0.05, **P□<0.01 ***P□<0.001 and ****P□<0.0001.

Other MAIT cell clusters contained cells from both species, but the representation of mouse versus human cells was highly unbalanced. For example, cells in *i*-cluster 3 expressed genes typical of MAIT1 cells, and were prevalent in mouse liver and spleen, but represented to a much lesser extent in human PBMCs **(Fig. 7C and Extended Data Fig. 9A)**. Integrated *i*-cluster 1 had a MAIT17 signature, including expression of *TMEM176A* and *IL18R1*, with mouse cells from different organs and only a few human cells, consistent with the prevalence of MAIT17 cells in mice **(Fig. 7C and Extended Data Fig. 9A)**. Mouse lung MAIT cells not only had a unique transcriptome compared to mouse MAIT cells from other sites, but human and mouse lung cells did not cluster together. Integrated *i*-cluster 2 consisted of lung mouse MAIT cells, but only a few human lung MAIT cells were present, demonstrating that the lung gene expression signature varies between the two species.

The divergent transcriptional signatures in peripheral MAIT cells might reflect genetic and/or environmental differences between the two species. To further understand how environment can affect the properties of MAIT cells, we have studied outbred mice from pet stores, so-called dirty mice, and neonatal SPF mice cross-fostered with pet store mothers. The number of lung MAIT cells was not greatly increased by exposure to the non-SPF flora (**Extended Fig. 9B, 9C**). We found alterations in MAIT cell phenotype that were environment-dependent, however, and likely related to the microflora. Pet store mice had fewer SCD1^+^ MAIT cells in the lung (**Fig. 7D**, **Extended Fig. 9B**), a marker of mature MAIT17 cells. The differences were more apparent, however, when cross-fostered C57BL/6 mice were included in the comparison, which eliminated a role for genetic differences (**Fig. 7D**, **Extended Fig. 9B**). Cross-fostered mice also had a reduction in SDC1**^+^** lung MAIT cells, but also had reduced expression of RORγt and increased percentages of KLRG1^+^ and T-bet^+^ lung MAIT cells, a sign of enhanced MAIT1 cell function (**Fig. 7D**, **Extended Fig. 9B**). These data suggest that removal from SPF conditions increases the prevalence of MAIT1 at the expense of MAIT17 cells.

## Discussion

Here we have used a combination of scRNA-seq, phenotypic and metabolic analyses to characterize the heterogeneity of MAIT cell subsets from different tissues and the conservation of these subsets between mouse and human. Our data revealed the presence of a subset of circulatory MAIT cells that in the mouse includes recent thymic emigrants. Furthermore, transcriptomic and measurement of metabolic parameters indicated that mature MAIT cells, even those in the thymus, had a metabolic state highly different from naïve CD8^+^ T cells, but only partially similar to subsets of memory CD8^+^ T cells in mouse and human. Additionally, the metabolic state of mouse MAIT1 and MAIT17 cells was strikingly different. While some subsets of mouse and human MAIT cells defined by scRNA-seq were well-represented and conserved between species, particularly in the thymus, other peripheral MAIT cell subpopulations had major quantitative and qualitative differences between mouse and human, that could reflect in part environmental influences.

In the thymus, we identified multiple subsets of mouse and human MAIT cells, including both precursors unique to the thymus as well as clusters of mature MAIT thymocytes that also were present in the periphery. When we analyzed the mouse thymus stages by single-cell trajectory analysis, we generated a model in which mouse precursors mature into stage 2-like MAIT cells and then diverge into distinct MAIT1 and MAIT17 subsets, with MAIT17 cells predominant. The finding of mature thymic MAIT1 and MAIT17 subsets agrees with several others^9, 10, 16, 21^, but here we also showed that maturation from stages 1-3 is accompanied by corresponding changes in metabolic parameters, with stage 3 cells more similar to peripheral MAIT cells. Unlike another report^22^, however, we did not find evidence for MAIT2 cells, which may reflect the difference between C57BL/6 and BALB/c strain mice. Our data suggest that MAIT cells with a stage 2-like phenotype, some that are CCR7^+^, also are found in the blood and spleen. Therefore, the data are consistent with a model in which partially mature MAIT cells are exported from the thymus, while other cells retained in the thymus attain a more mature or stage 3 state, similar to a current model for iNKT cell differentiation^42^. We speculate that it provides a strong advantage for the host to have MAIT subsets that can readily differentiate on-demand in the periphery in response to an infection.

The evidence indicates human thymus MAIT cells differentiate from precursors mostly into a MAIT1 cell population. In agreement with an earlier study^10^, an early MAIT cell subset (human cluster 6) with cells expressing *RAG1* and *RAG2* found in humans was absent in the mouse thymus. This might reflect differences in the kinetics of differentiation, for example, the transition from the most immature stage (human cluster 6) might be slower in the human compared to the mouse thymus. In addition to progressing into MAIT1 cells, the pseudotime analysis suggests some human MAIT thymus precursors differentiate to cells that expressed CCR7 and have other features similar to mouse stage 2 MAIT thymocytes (Fig. 5). The analysis could not determine, however, if these cells differentiate further. Additionally, a population of human thymus MAIT17 cells was not detected.

Overall, our data show that populations of mouse and human MAIT cells have metabolic features that distinguish them from naïve CD8^+^ T lymphocytes. The predominant mouse MAIT17 cell subset had a highly active metabolism characterized by fatty acid uptake and storage and mitochondrial activity, while MAIT1 cells were more active in glucose uptake. This metabolic difference was correlated with their function and was observed across tissues, including in the thymus. Similar findings were recently reported when γδ T cells that produce IFNγ were compared to those that produce IL-17, highlighting similarities t between populations of innate-like T lymphocytes^51^. As for the transcriptomes, subsets of human MAIT cells from lung and blood were not greatly different from one another, nor could they be divided into subgroups by surface phenotype. Instead, the human MAIT cell subsets shared similar metabolic features, most comparable to tissue-resident CD8^+^ memory T cells for uptake and storage of fatty acids but with a reduced mitochondrial potential. Interestingly, this metabolic phenotype resembles the controlled activation state of epithelial-resident T cells^52^ A previous study also showed generally low mitochondrial activity in bulk MAIT cells from PBMCs, with an ability to rapidly reactivate metabolic and effector pathways upon stimulation^53^. Increased mitochondrial potential and activity was functionally linked to increased IL-17 production by human MAIT cells^54^ consistent with a connection between mitochondrial activity, IL-17 production by MAIT cells and the different metabolic state of mouse MAIT17 cells.

Increased production of IL-17 by human MAIT cells has been observed in several contexts, including children with community-acquired pneumonia^27^. In multiple studies, however, the proportion that produced IL-17 was much lower than the frequency of those producing IFNγ, and it is uncertain if there is a true MAIT17 subset, as opposed to more flexible or polyfunctional cells that also had the capability to produce IFNγ. Some human MAIT cells with an IL-17 gene expression signature were present in lung, but the MAIT1 gene expression signature was present in MAIT cells in these clusters as well. All considered, including the data from the thymus, we conclude that an intrinsic, highly specialized MAIT17 subset either is absent or very rare in humans.

There is evidence that obesity and metabolic alterations can alter MAIT cell function. This has not only been observed in mice^55^, but also there was increased MAIT cell production of IL-17 by MAIT cells from obese individuals^54, 56^, although IFNγ-producing cells remained more numerous. Furthermore, supplementation with the TCA metabolite alpha-ketoglutarate augmented human MAIT cell effector capacity^56^ providing a further connection between metabolism and MAIT cell function. These data suggest there is a causal link between metabolism and MAIT cell function, although further studies will be required to establish this.

MAIT cell specificity is highly conserved and therefore we examined the extent to which the transcriptomes of MAIT cells were also conserved. The data reveal the most similarity between differentiating human and mouse MAIT cells in the thymus. Even in peripheral MAIT cells, some homologous genes were regulated similarly in the two species, especially for MAIT1 cells. Furthermore, strong tissue differences were observed. For example, lung MAIT cells in both species were different from their counterparts in other tissues. Mouse and human lung MAIT cells did not align well, however, in the integration analysis, nor did the transcriptome of mouse spleen MAIT1 cells align with human MAIT1 cells from PMBC.

Undoubtedly, genetic differences between mice and humans influence the frequency and function of the MAIT cell population, but it is also possible that the highly controlled, standard SPF conditions of laboratory mouse housing have an influence as well. Exposure to the intestinal microbiome is not only necessary for MAIT cell thymic development^57^, but also differences in the microbiome can influence the number and function of skin MAIT cells^57^. Our data indicate that exposure to a less controlled environment increased cells with a MAIT1 phenotype and decreased MAIT17 cells. It remains to be determined the extent to which differences between mouse and human MAIT cell transcriptomes, and ultimately function, can be attributed to differences in microbial exposure as opposed to other environmental factors and genetic differences.

## MATERIALS AND METHODS

### Animals

Inbred mice were bred and housed under specific pathogen-free conditions in the vivarium of the La Jolla Institute for Immunology (La Jolla, CA). C57BL/6J mice were purchased from Jackson laboratories. Rag2:GFP mice were obtained from Dr. Pamela Fink at University of Washington. Female mice were used and they were 6–12 weeks old, unless indicated otherwise. Pet shop mice were analyzed immediately after purchase or housed in a vivarium maintained by the University of California, San Diego. We used SPF (specific pathogen free) C57BL/6 mice of similar weight as controls because the precise age of pet store mice was unknown. For cross-fostering, breeding pairs of SPF B6 mice and pet shop mice were simultaneously set up when individual mice reached approximately 6 weeks of age. SPF B6 pups born within 48 h were used for cross-fostering. After the birth of both SPF B6 and pet shot litters, the pet shop litters were removed and replaced with similar numbers of pups from the SPF B6 litters. Litters from SPF B6 breeders were then nursed by pet shop mothers until weaning. Cross-fostered male and female mice were analyzed when they were approximately 8 weeks of age. All procedures were approved by the La Jolla Institute for Immunology or University of California San Diego Animal Care and Use Committee and are compliant with the ARRIVE standards.

### Antibodies and tetramers

Mouse and human MR1 tetramers loaded with either 5-OP-RU or 6-FP were obtained from the NIH Tetramer Core Facility. Fluorochrome-conjugated monoclonal antibodies were purchased from eBioscience, BD Bioscience, or BioLegend. Antibodies with clone indicated in parentheses: anti-mouse CD45 (30-F11); anti-mouse IgD (clone 11-26c.2a); anti-mouse γδ TCR (clone GL3); anti-mouse CD4 (clone GK1.5 or RM4-5); anti-mouse CD8α (clone 53-6.7); anti-mouse CD8β (clone H35-17.2); anti-mouse CD138 (clone 281-2); anti-mouse TOX (clone TXRX10); anti-mouse CD19 (clone 1D3); anti-mouse CCR9 (clone CW-1.2); anti-mouse CD24 (clone M1/69); rabbit polyclonal anti-mouse LEF1 (C12A5); anti-mouse SATB1 (clone 14/SATB1); anti-mouse IFN-γ (clone XMG1.2); anti-mouse TNF (clone MP6-XT22); anti-mouse IL-17A (clone TC11-18H10); anti-mouse CD69 (clone H1.2F3); anti-mouse T-bet (clone O4-46); anti-mouse RORγT (clone Q31-378 or B2D); anti-mouse IL-18R1 (clone BG/IL18Ra); anti-mouse CCR7 (clone 4B12); anti-mouse CD11b (clone. M1/70); anti-mouse CD62L (clone MEL-14); anti-mouse CD45R/B220 (clone RA3-6B2); anti-mouse CD11c (clone N418); anti-mouse ICOS (clone C398.4A); anti-mouse CXCR3 (clone CXCR3-173); anti-mouse TCRβ (clone H57-597), anti-mouse CD44 (clone MI7); anti-mouse KLRG1 (clone 2F1); anti-human CD3 (clone OKT3); anti-human CD8 (clone RPA-T8); anti-human CD161 (clone HP-3G10); anti-human Vα7.2 (clone 3C10); anti-human CD19 (clone HIB19); anti-human CCR7 (clone 150503); anti-human CD45RA (clone HI100) and anti-human CD103 (clone Ber-ACT8).

### Isolation of mouse cells

Splenocytes and thymocytes were harvested by mechanical disruption on 70 μm cell strainers followed by red blood cell (RBC) lysis and washing with Hank’s Balanced Salt Solution (HBSS) (Gibco) supplemented with 10% FBS. Lung tissue was digested with STEMCELL spleen dissociation medium, and mechanically dissociated using GentleMACS Dissociator (Miltenyi). Cells were strained though a 70 μm filter and washed with HBSS supplemented with 10% FBS followed by RBC lysis. Liver cells were harvested by mechanical disruption on 70 μm cell strainer followed by 34% Percoll gradient before RBC lysis and washing.

### Flow cytometry

For staining of cell surface molecules, cells were suspended in staining buffer (PBS, 1% bovine serum albumin (BSA), and 0.01% NaN_3_) and first stained with using PE- or APC-conjugated MR1 tetramers at a dilution of 1:300 in staining buffer for 45 minutes at room temperature followed by surface staining with fluorochrome-conjugated antibody at 0.1–1 μg/10^7^ cells. Cells were stained with Live/Dead Yellow (ThermoFisher) at 1:500 and Fc receptors were blocked with 2.4G2 antibody at 1:500 and Free Streptavidin at 1:1000 for 15 min at 4°C. After washing, cells were stained with cell surface-specific antibodies for 30 minutes on ice. For cytokine staining, cells were previously stimulated with 100 ng/ml of PMA and 1 μg/ml of Ionomycin for 1h at 37°C and then incubated in GolgiStop and GolgiPlug (both from BD PharMingen) for 2 h at 37°C. For intracellular staining, cells were fixed with CytoFix (BD) for 20 min, and permeabilized with Perm 1X solution (ThermoFisher) with intracellular antibodies overnight. For high-parameter flow cytometry experiments, data were acquired on Fortessa or Symphony S6 (BD Biosciences), data were processed with DIVA (BD Bioscience) and analyzed with FlowJo v10.7 (BD). Opt-tSNE and UMAP dimensional reduction as well as FlowSOM algorithm clustering of flow cytometry data was performed in OMIQ software (OMIQ Inc.).

### Cell enrichment and cell sorting

For MAIT cell enrichment before sorting, negative selection of cells was carried out using biotinylated antibodies against CD11b (clone M1/70), CD11c (clone M418), F4/80 (clone BM8.1), CD19 (clone 1D3), and TER-119 (clone TER-119). These were used together with Rapidspheres (STEMCELL Technologies) and either the Big Easy (STEMCELL Technologies) or Easy eight magnets (STEMCELL Technologies) and protocols from Stem Cell Technologies. MAIT cells were sorted using a FACSAria III (BD Biosciences).

### Human tissue and cell preparation

Postnatal human thymus was obtained from children with congenital heart disease undergoing cardiac surgery at Rady Children’s Hospital, San Diego, CA. Only patients who meet the inclusion criteria and sign informed consent, are included in the study. Thymus samples are obtained from 2-year-old male, 2-year-old female, two 13-month-old males and 4-year-old female. Thymus tissue was processed by mechanical dissociation into a single cell suspension, strained and viable lymphocytes were purified by Lymphoprep (STEMCELL) density gradient centrifugation before cryopreservation. For lung and peripheral blood samples, written, informed consent was obtained from all subjects from the Institutional Review Board of La Jolla Institute for Immunology and the Southampton and South West Hampshire Research Ethics Board. Newly diagnosed, untreated patients with non-small cell lung cancer were prospectively recruited once referred. Freshly resected tumor tissue and, where available, matched adjacent non-tumor lung tissue was obtained from patients with lung cancer following surgical resection. Lung tissue was obtained from 76-year-old female, 63-year-old female, 66-year-old male and 48-year-old male. Tissues were macroscopically dissected and slowly frozen in 90% FBS (Thermo Fisher Scientific) and 10% DMSO (Sigma) for storage, until samples could be prepared. Cryopreserved non-tumor lung tissue was mechanically dissociated and enzymatically digested as previously described^45^. Briefly, lung tissue was minced with a scalpel and digested enzymatically with 0.15 WU/mL of D-Liberase (Roche) and 800□U/mL of DNase I (Sigma-Aldrich) for 15□min at 37°C. Then it was disaggregated into a single-cell suspension by passing it through a 70□µm strainer and rinsing with cold buffer (1× phosphate-buffered saline (PBS), 2□mM EDTA, 0.5% BSA). PBMCs are either obtained from same patients we obtained lung from or from healthy donors. PBMCs were isolated using density gradient before cryopreservation.

### Metabolic assays

Cytometry-based metabolic assays have been described previously^58^. Briefly, cells were stained with MitoTracker Deep-Red FM (Life Technologies) at 100 nM concentration, 37 °C, 5 % CO_2_ for 30-45 minutes in RPMI1640 (Gibco) containing 5 % FBS. For glucose uptake measurements, cells were incubated in glucose-free media containing 5 μg/ml 2-(N-(7-Nitrobenz-2-oxa-1,3-diazol-4-yl)Amino)-2-Deoxyglucose (2-NBDG, Thermo Fisher) and 2.5% FBS at 37 °C, 5 % CO_2_ for 30 minutes, unless indicated otherwise. For lipid droplet quantification, cells were incubated in media containing 1 µg/ml Bodipy 493/503 (Thermo Fisher) for 30 min. Uptake of fatty acids was quantified after incubation with 1uM 4,4-Difluoro-5,7-Dimethyl-4-Bora-3a,4a-Diaza-s-Indacene-3-Hexadecanoic acid (Bodipy-FL C16, Thermo Fisher) at 37 °C, 5 % CO_2_ for 30 minutes. Optimal incubation periods for metabolic dye and metabolite uptake depend on the tissue and required fluorescence intensity, but only exceeded 45 minutes where indicated. Data were acquired using Fortessa or LSR II flow cytometers (BD Biosciences) and analyzed with FlowJo v10.7 software (BD Life Sciences). Metabolic marker fluorescence intensity depends on the instrument type and laser intensity, and therefore does not allow inter-experiment comparisons.

### Single-cell RNA sequencing

Cells were sorted into a low retention 1.5 mL collection tubes containing 500 μL of a solution of PBS: FBS (1:1) supplemented with RNase inhibitor (1:100). After sorting, ice cold PBS was added to make up to a volume of 1.4 mL. Cells were then spun down (5 min, 600 *g*, 4°C) and the supernatant was carefully aspirated, leaving 5 to 10 μl. The cell pellet was gently resuspended in 25 μl of resuspension buffer (0.22 μm filtered ice cold PBS supplemented with ultra-pure BSA; 0.04%, Sigma-Aldrich). Following that, 33 μl of the cell suspension was transferred to a PCR-tube and single-cell libraries prepared as per the manufacturer’s instructions (10x Genomics). Samples were processed using 10X v2 chemistry for the mouse dataset and 10X v3 chemistry for the human dataset, as per the manufacturer’s recommendations; 11 and 12 cycles were used for cDNA amplification and library preparation, respectively. Libraries were quantified and pooled according to equivalent molar concentrations and sequenced on Illumina HiSeq 2500 or NovaSeq sequencing platform with the following read lengths: read 1 – 101 cycles; read 2 – 101 cycles; and i7 index - 8 cycles.

### Single-cell transcriptome analysis

Mouse cell libraries were mapped with Cell Ranger’s count pipeline for mm10. Then multiple libraries were aggregated with the aggr pipeline. Aggregated data were then imported into the R environment where Seurat^59^ (2.1.0) was used to filter and find clusters. Cells with less than 200 genes and more than 2,500 genes were discarded. Furthermore, cells with more than 5% UMIs coming from mitochondrial genes were filtered out. Genes expressed in less than 3 cells were ignored. This resulted in 6,080 cells with 13,503 genes for downstream analyses. The gene expression matrix was then normalized and scaled. Principal Component Analysis was performed on the scaled data and, based on the elbow plot, 20 principal components were selected for clustering, default resolution (0.6) was used and a perplexity of 100 was chosen for the t-SNE dimensionality reduction. This dataset was further split up into 3 tissue types – spleen (3145 cells), lung (1313 cells), thymus (535 cells) which were then analyzed individually using the same steps. For lung - 8 PCs; thymus – 21 PCs and for spleen – 15 PCs were used for clustering.

Human cell libraries were mapped with Cell Ranger’s count pipeline for reference GRCh38-1.2.0. Multiple libraries were aggregated with the aggr pipeline. Aggregated data was then imported into the R environment where Seurat (v3.9.9.9008) was used to filter cells, normalize and find clusters. Cells with less than 200 genes and more than 20,000 UMIs were discarded. Furthermore, cells with more than 15% UMIs coming from mitochondrial genes were filtered out. Genes expressed in less than 3 cells were ignored. This resulted in 3,020 cells with 17,626 genes for downstream analyses. The gene expression matrix was then normalized and scaled using log normalization. Principal Component Analysis was performed on the scaled data, and based on the elbow plot, 18 principal components were selected for clustering, default resolution (0.9) was used. To determine the clusters’ enriched genes (markers), Seurat’s FindAllMarkers function was used with test.use = MAST (Adjusted P-value < 0.05 and |log fold change| > 0.25). For analyzing human thymus, cells (1316) from thymus tissue type were selected and analyzed using the same steps as listed above. 25 PCs were used for clustering of the thymus cells.

### Human-mouse data integration

Single-cell sequencing data from human and mouse MAIT cells was integrated using Seurat’s (3.0.2) alignment method^50^. Briefly, we identified cross-dataset pairs of cells matching biological states (anchors). These pairs are used to correct technical differences between conditions. Default parameters were used and the following was tailored: the first 15 principal components based on the elbow plot; and resolution 0.5 was used to identify the clusters in the integrated data.

### Signature plots

Signature module scores were calculated with Seurat’s AddModuleScore function using default parameters. This function calculates the average expression levels of a gene set of interest, subtracted by the aggregated expression of control gene sets, randomly selected from genes binned by average expression Gene lists used for MAIT1 and MAIT17 analysis were obtained from Legoux et al, 2019^21^. Gene lists for tissue resident memory and circulating signatures were obtained from Milner et al., 2017^39^. Gene lists for glycolysis was obtained from MSigDB geneset KEGG_GLYCOLYSIS_GLUCONEOGENESIS. Gene list for oxidative phosphorylation was obtained from MSigDB geneset HALLMARK_OXIDATIVE_PHOSPHORYLATION. Gene list for Fatty acid metabolism was obtained from MSigDB geneset KEGG_FATTY_ACID_METABOLISM. Gene list for mitochondrial gene was obtained from MSigDB geneset MITOCHONDRIAL_GENE_EXPRESSION.

### Single-cell trajectory analysis

A wrapper script from R package SeuratWrappers v0.3.0 was used for Calculating Trajectories with Monocle3^60^ v0.2.3.0. Seurat’s object with Seurat clustering and UMAP coordinates was converted into a Monocle3 object. A single monocle partition was used for all cells. Monocle3 function learn_graph was used to fit a principal graph for the partition used. The cells were then ordered using the function order_cells, which calculates where each cell falls in pseudotime. Monocle3 helper function get_earliest_principal_node was used to specify the root node of the trajectory. Trajectory UMAPs were plotted using the function plot_cells with Monocle3 object as input. The function graph_test was used to find genes that are differentially expressed on different paths through the trajectory with the option neighbor_graph=“principal_graph”. The trajectory-variable genes were then collected into co-regulated modules using the function find_gene_modules. Monocle3’s function aggregate_gene_expression was used to calculate aggregate expression of genes in each module for all the clusters. These module scores were then plotted in a heatmap using R package pheatmap v1.0.12 with options cluster_rows=TRUE, cluster_cols=TRUE, scale=“column”, clustering_method=“ward.D2”. The modules were further combined into stages based on their functionality/annotation. Top 10 genes were selected for each stage with high morans_I value calculated by the function graph_test earlier. The Module heatmap was generated using Seurat’s function DoHeatmap. It shows scaled average expression for the top 10 genes grouped by cells in each stage.

### Demultiplexing

PLINK (v2.1.4) from Illumina Genome Studio plugins was used to convert and export Illumina genotype data into PLINK data format. PLINK is again used with the “--recode vcf” option to convert PLINK data format to VCF. snpQC package^61^ was used to detect low quality SNPs. SNPs failing in >5% of the samples and SNPs with Illumina’s gene call scores <0.2 in <90% of the samples were excluded for downstream analysis. The alignment files generated by CellRanger count program (v3.1.0) were split by cell barcode using samtools (v1.9)^62^. Each individual cell-specific BAM file was run through Freebayes (v0.9.21)^63^ with the SNP array variants as input to catalog matching SNPs for each cell.

The score for each sample-cell barcode pair were calculated as shown in the below formula:

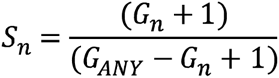

where G_n_ is the number of genotype calls for this barcode that match sample n and G_ANY_ is the total number of genotype calls made for this barcode. Scores for each sample were then ranked from highest to lowest and the score for the highest-ranking sample was compared to that of the second highest. If the ratio between these two was 1.3 or greater and at least 300 genotype calls were made for the cell, the sample with the highest score was assigned.

### Data analysis using Compass algorithm

Cellular metabolic states were inferred from single cell transcriptomic data and flux balance analysis using the Compass algorithm^43^ v0.9.9.6.3. Briefly, Compass algorithm was fed with the count matrix from MAIT17 (cluster 0, 5 and 9), MAIT1 (cluster 1), circulatory (cluster 2, 4 and 7), lung tissue resident (cluster 3) and precursor (cluster 6) MAIT cells from the mouse scRNA-seq data. This matrix was obtained after performing single-cell analysis using Seurat. No cell during the Compass analysis was aggregated into microclusters. The resulting “reaction scores matrix” output was then subjected to downstream analyses. We selected the reactions labeled by Recon2^44^ GSMM as involved in “Glycolysis/gluconeogenesis”, “Citric acid cycle”, and “Fatty acid oxidation” pathways, and for Amino acid metabolism, we used the reactions filtered by Compass developers. For all the reactions, we included only the ones whose Recon2 confidence is either 0 or 4 and are annotated with an EC (Enzyme Commission) number, according to the reaction metadata included in the Recon2 database. We kept reactions with unevaluated confidence (Recon2 confidence score of 0) because some of them were found to be key reactions in primary metabolic pathways, but excluded the ones that Recon2 curators explicitly specified to not have direct biochemical support (Recon2 confidence score of 1, 2 and 3), according to Compass developers. To find reactions with differential potential activity based on Compass predictions, we computed the analysis of variance (ANOVA) for each reaction of the Compass scores matrix. The resulting p-values are adjusted with the Benjamini-Hochberg (BH) method and were added as a new column value as “adjusted p-value”. We defined a reaction as significantly differentially expressed if the adjusted p-value is smaller than 0.1, same as the Compass developers. Effect size was further assessed with Cohen’s D statistic, which is defined as the difference between the sample means over the pooled sample standard deviation.

Let n_1_, x_1_, s_1_ be the number of observations in population 1, the sample mean and standard deviation of their scores in a given reaction, respectively. (With a similar notation for population 2). Then

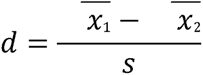

with

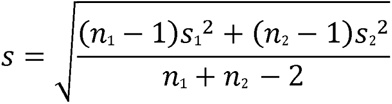

Paired effect size analysis was performed for different MAIT cell subsets as compared to precursors of mature MAIT cells. The equation was applied for each paired comparison. Taking one of the different MAIT cell subsets as population 1 and the precursors of mature MAIT cells as population 2.

### Statistical Methods

All graphs and statistical analysis were generated using Prism 9 software (GraphPad Software, San Diego, CA). Data are plotted as mean ± standard deviation or mean ± standard error of the mean (SEM), and statistical significance was determined by using unpaired t test. Significance for multiparameter comparisons was determined by one-way ANOVA with Dunnett’s post-test, paired t test or one-way ANOVA with post-hoc Tukey test.

### Data visualization tool

Visualization web-based platform was constructed using R package Shiny v1.7.1, customizing it with CSS theme and htmlwidgets. The app was fed with mouse and human Seurat and Monocle objects, subsetting only the elementary data in order to optimize memory resources. Plots are constructed in real time using Seurat and Monocle V3 functions using the single cell data object selected by the user.

### Data and Software Availability

Single-cell RNA sequencing data generated for this study are deposited at the Gene Expression Omnibus under accession number GSE189485. Data visualization tool is available at https://mait.lji.org

### Code availability

The code developed for the analyses performed in this study is available upon request.

## Acknowledgement

We would like to thank Anusha Preethi Ganesan for help with human lung dissociation protocol and Gina Levi for coordinating receipt of human thymus samples. We thank the La Jolla Institute (LJI) Flow Cytometry Core for assisting with cell sorting and sequencing Core for performing scRNA sequencing. Supported by the US National Institutes of Health grants AI105215, AI71922 and AI137230 to M.K., Wellcome Trust grant 210842_Z_18_Z to T.R, NIH grants AI108651 and AI163813 (L.-F.L) and UCSD Program in Immunology seed grant (L.-F.L and S.M.H). Lung tissue collection in the UK was supported by funding from the Whittaker fund and iCURE, by the Wessex Clinical Research Network and the National Institute of Health Research, UK. We thank Mr. Woo and Mr. Alzetani for access to surgical material, Dr. Serena Chee, Ben Johnson, Alice Appleford and Sophie Matthew for collection, processing and storage of the tissues.

Utilized equipment was supported by the NIH grants no. S10RR027366 (BD FACSAria II), S10OD025052(NovaSeq 6000) and no. S10-OD016262 (Illumina HiSeq 2500).

## Author contributions

S.C., G.A, T. R., S.M.H., L.-F.L., and M.K. designed the experiments, which were performed by S.C., G.A, T. R., G.S., H.S., M.P.M., G.Y.S. and C.H.L. Data analysis was performed by S.C., G.A, T. R., A.S., C.R., V.C.C. A.L.R.P. and J.G. Human thymus samples were provided by J.L., R.M. and J.N. in collaboration with H.C. Thymus samples were processed by G.V. and Y.L. Human lung samples were provided by C.O. in collaboration with P.V. Manuscript was written by S.C., G.A., T.R. and M.K. with editing by H.C., C.O., S.M.H., L.-F.L., and P.V. All authors reviewed the manuscript.

---

**Extended Data Figure 1:**
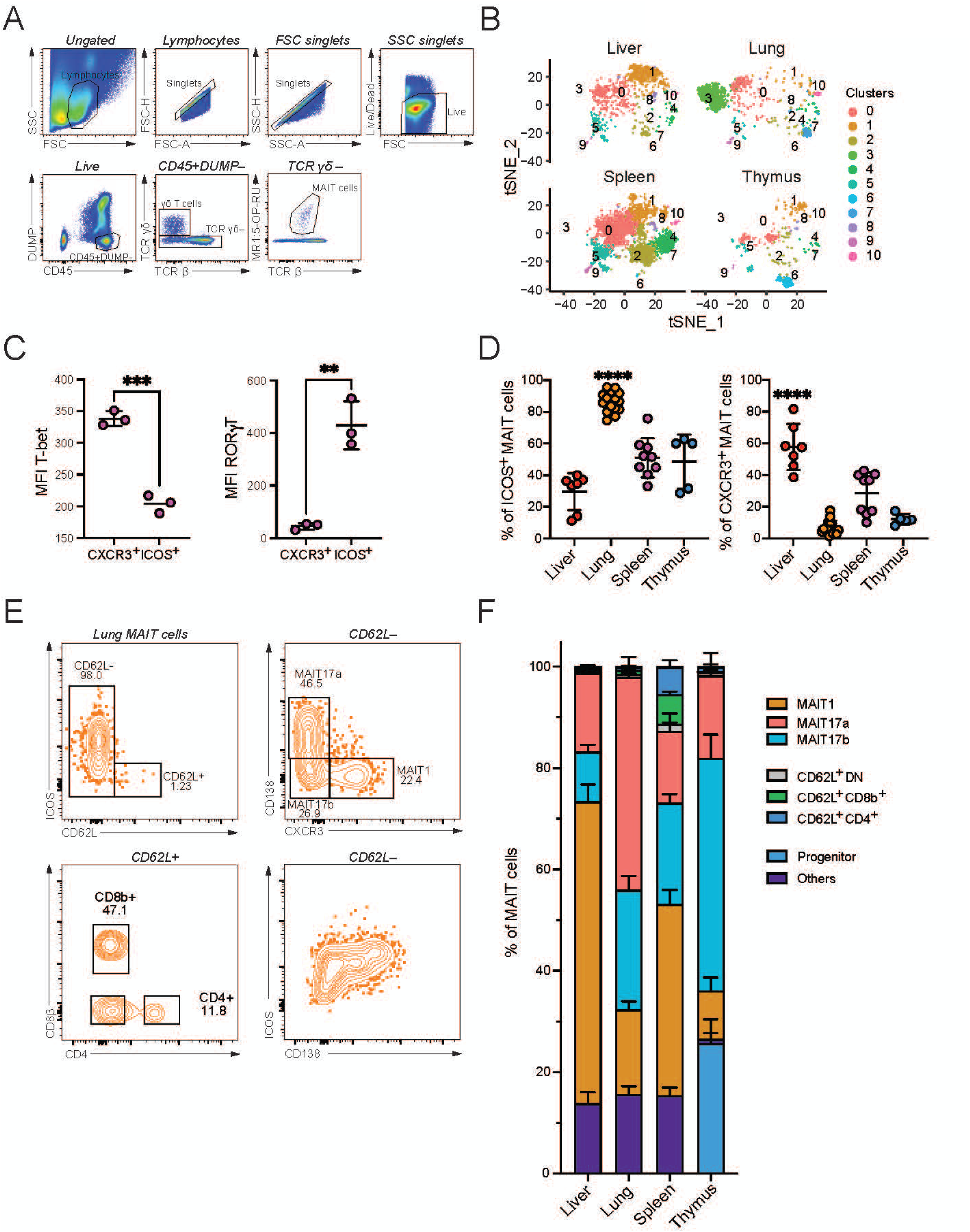
Characterization of mouse MAIT cell subsets. (A) Representative flow cytometry gating used to identify MAIT cells in different tissues. Live/Dead Yellow negative single cell events were gated by excluding antigen-presenting cells (DUMP: CD11c, CD11b, IgD, B220) and γδ T cells from the CD45^+^ population, CD4^+^, CD8^+^ double positive cells were excluded as well. (B) t-SNE plot showing the degree to which MAIT cell clusters are composed of cells from the indicated mouse tissues. Each cluster has the same t-SNE coordinates as in Fig. 1A. (C) MFI indicating expression of T-bet and RORγT transcription factors in spleen MAIT cell subpopulations defined by surface markers as MAIT1 (CXCR3^+^) and MAIT17 (ICOS^+^). Paired t-test. n = 3. *: p < 0.05; **: p < 0.01; (D) Frequency of CXCR3^+^ and ICOS^+^ MAIT cells in the indicated tissues. One-way ANOVA with post-hoc Tukey test. Liver n = 7, Lung n= 15, Spleen n= 9, Thymus n= 5. ****: adjusted p < 0.0001. (E) Lung MAIT cell subpopulations were detected by flow cytometry according to markers for MAIT cell clusters determined by scRNA-seq. The total MAIT cell gate was initially separated by CD62L expression. The CD62L negative gate was further divided into MAIT17a (ICOS^+^CD138^+^CXCR3^-^), MAIT17b (ICOS^+^CD138^-^CXCR3^-^) or MAIT1 (ICOS^-^ CD138^-^CXCR3^+^). The CD62L positive gate was divided into CD4^+^, CD8β^+^ or double negative (DN) (CD4^-^CD8b^-^CD62L^+^) subpopulations. (F) Percentage of each MAIT cell subpopulation, as defined above, in different tissues. Using the global gating strategy defined in (E) in combination with thymus-specific markers to detect immature MAIT cells (CD44^-^CD24^+^CCR9^+^LEF1^+^SATB1^+^), the proportion of MAIT cell subpopulations was determined for each tissue of 11 female C57BL/6 mice (12.3±6.1 weeks-old). Data from 5 additional thymus tissues also were included.

**Extended Data Figure 2:**
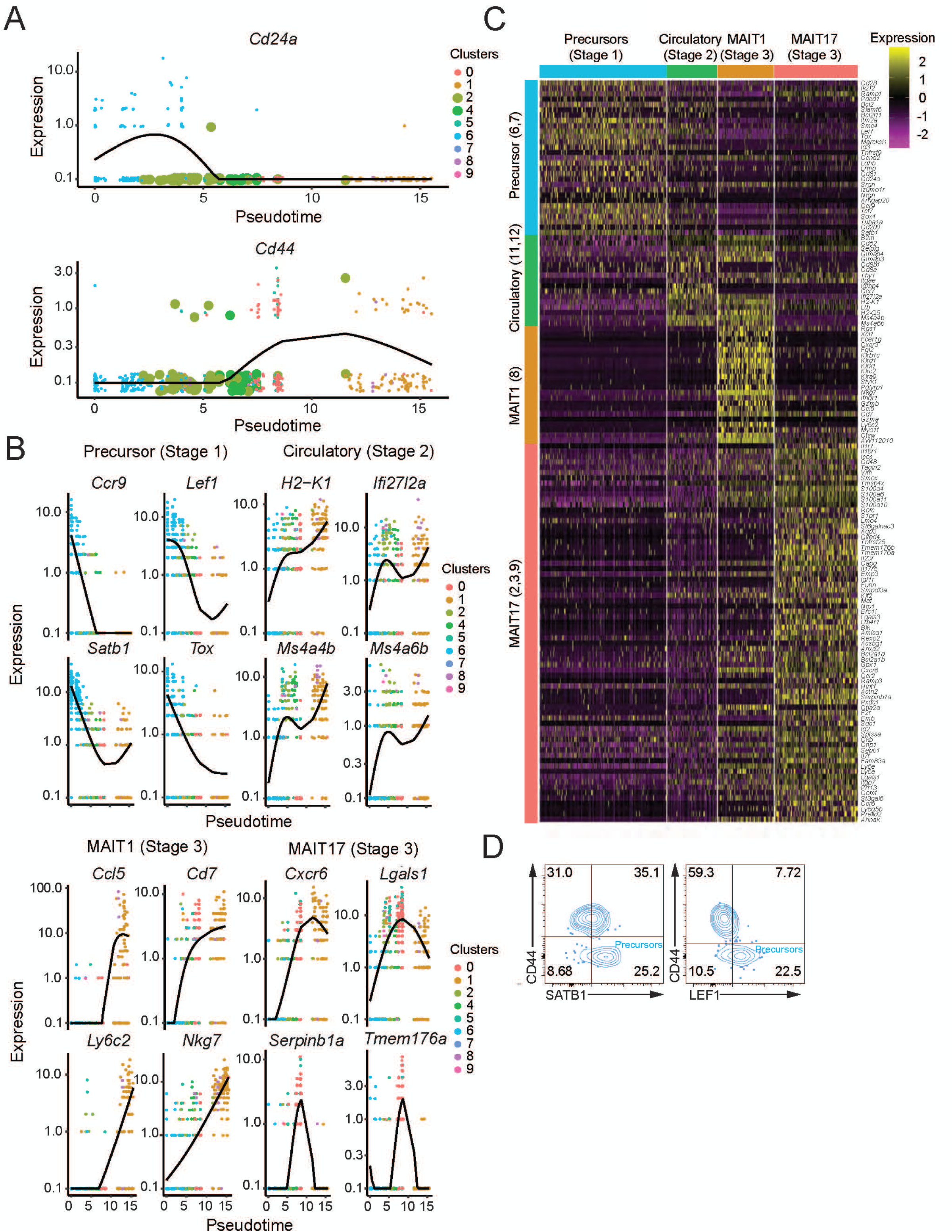
Transcriptional signatures reveal different stages of thymus MAIT cell differentiation. (A) Expression of *Cd24a* and *Cd44* along the pseudotime trajectory for MAIT thymus cells as constructed by Monocle 3. (B) Expression of the indicated stage-specific genes along the pseudotime trajectory as constructed by Monocle 3. (C) Scaled average expression heatmap of all the significantly differentially expressed genes along the MAIT thymus trajectory with Morans_I >0.2. Heatmap shows cells from the indicated MAIT cell differentiation stage and clusters from scRNA-seq on the *x*-axis. The gene modules to which the stage-specific genes belong are shown on the *y*-axis. (D) Representative flow cytometry plots for staining of gated, thymus MAIT cells for intracellular SATB1 or LEF1 along with surface expression of CD44, n = 5, from 2 experiments.

**Extended Data Figure 3:**
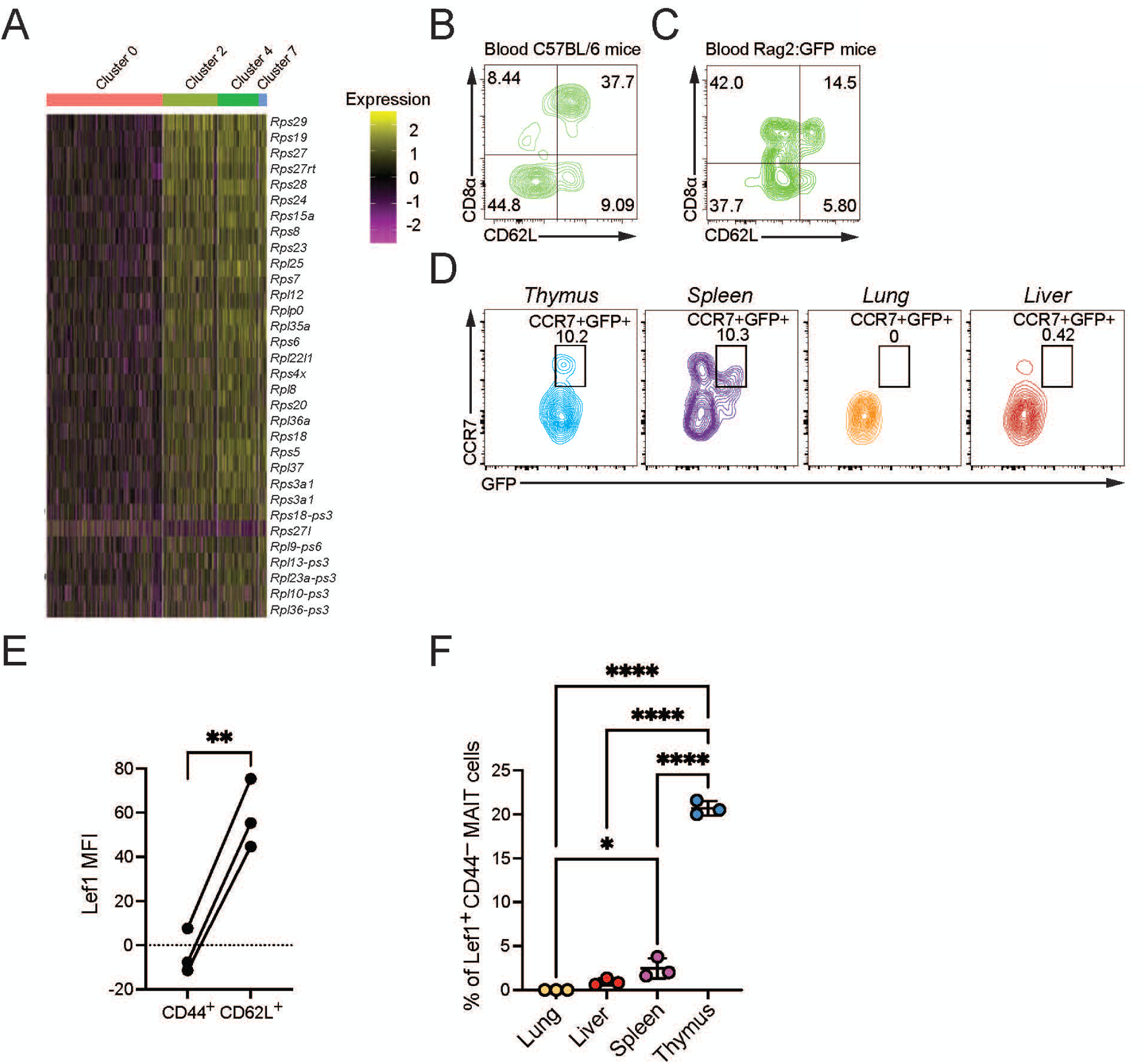
Circulatory MAIT cells. (A) Heatmap showing expression of the 25 most significantly enriched transcripts encoding ribosomal proteins in clusters 2, 4 and 7 in comparison with cluster 0. Representative flow cytometry plots for the expression of CD62L and CD8α in blood MAIT cells from C57BL/6 mice (B) and Rag2:GFP mice (C). (D) Representative expression of CCR7 and GFP in Rag2:GFP reporter mice. The percentage of MAIT cells that are CCR7^+^, GFP^+^ is indicated. (E) Intracellular staining for the transcription factor LEF1 in CD44^+^ or CD62L^+^spleen MAIT cells. Paired t-test, **: p-value < 0.01. (F) Percentage of LEF1^+^ CD44^-^ MAIT cells in different tissues. Intracellular staining for LEF1 in MAIT cells from liver, lung, spleen and thymus. One-way ANOVA with post-hoc Tukey test. n = 4. ****: adjusted p-value < 0.0001., (E, F) 16 week-old female mice.

**Extended Data Figure 4:**
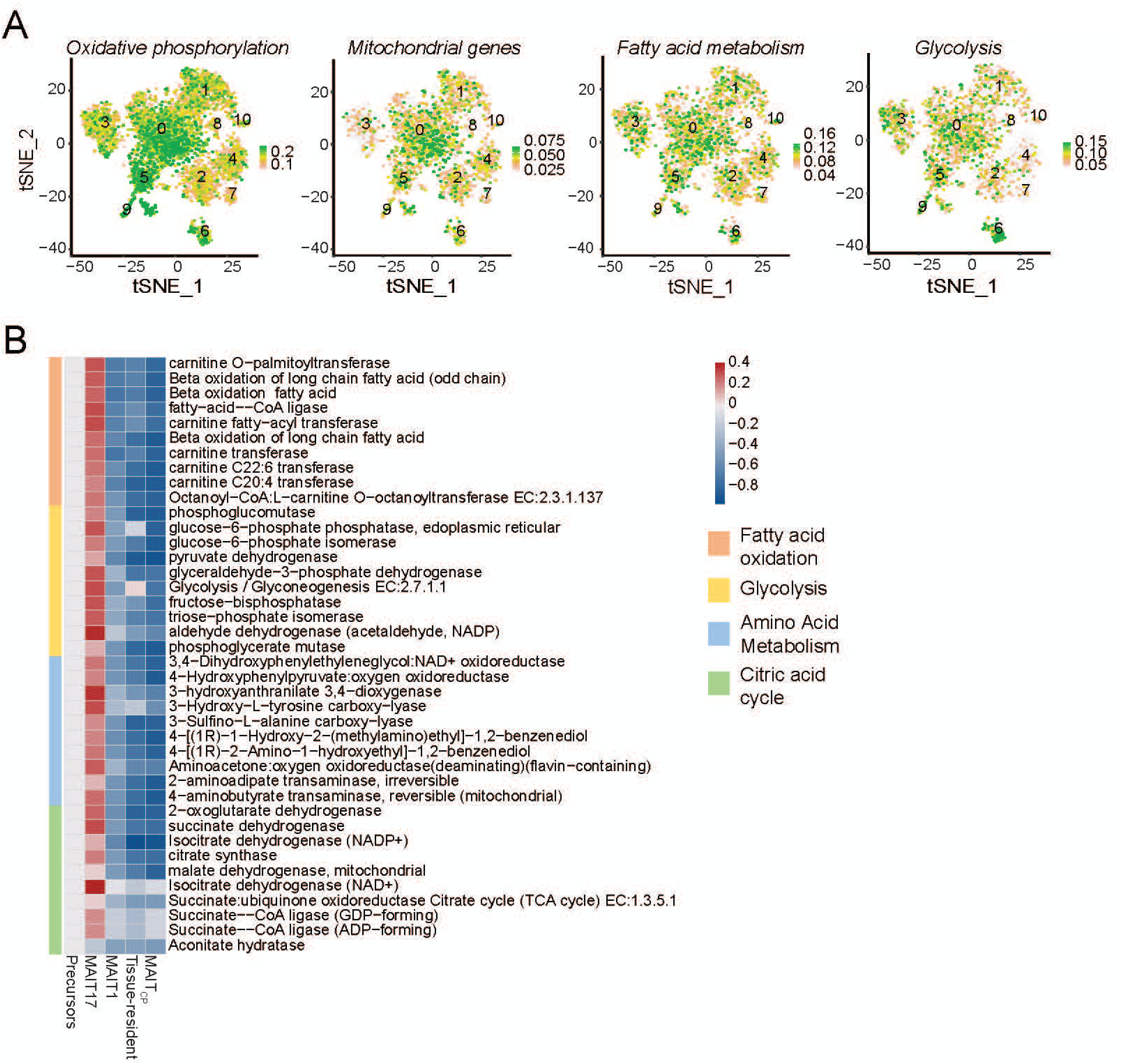
Different metabolic signatures of MAIT cell subsets. (A) t-SNE showing oxidative phosphorylation, mitochondrial gene, fatty acid metabolism and glycolysis signature scores for MAIT cells from the four sites. (B) Compass algorithm was used to assess the metabolic heterogeneity of MAIT cells. Progenitor = cluster 6; MAIT17= cells from clusters 0, 5 and 9; MAIT1= cluster 1; lung tissue resident= cluster 3; circulatory= clusters 2, 4 and 7). All the subsets of MAIT cells were compared to progenitor cells. Effect size with Cohen’s d statistic was calculated between each subset of MAIT cell in comparison to progenitor cells. Cohen’s d values were used for the color scale to represent in which MAIT subset each reaction pathway is being more or less (red or blue, respectively) active as compared to the progenitor cluster 6 cells. Top 10 genes are shown for fatty acid oxidation, glycolysis, amino acid metabolism and citric acid cycle with the lowest adjusted p-values.

**Extended Data Figure 5:**
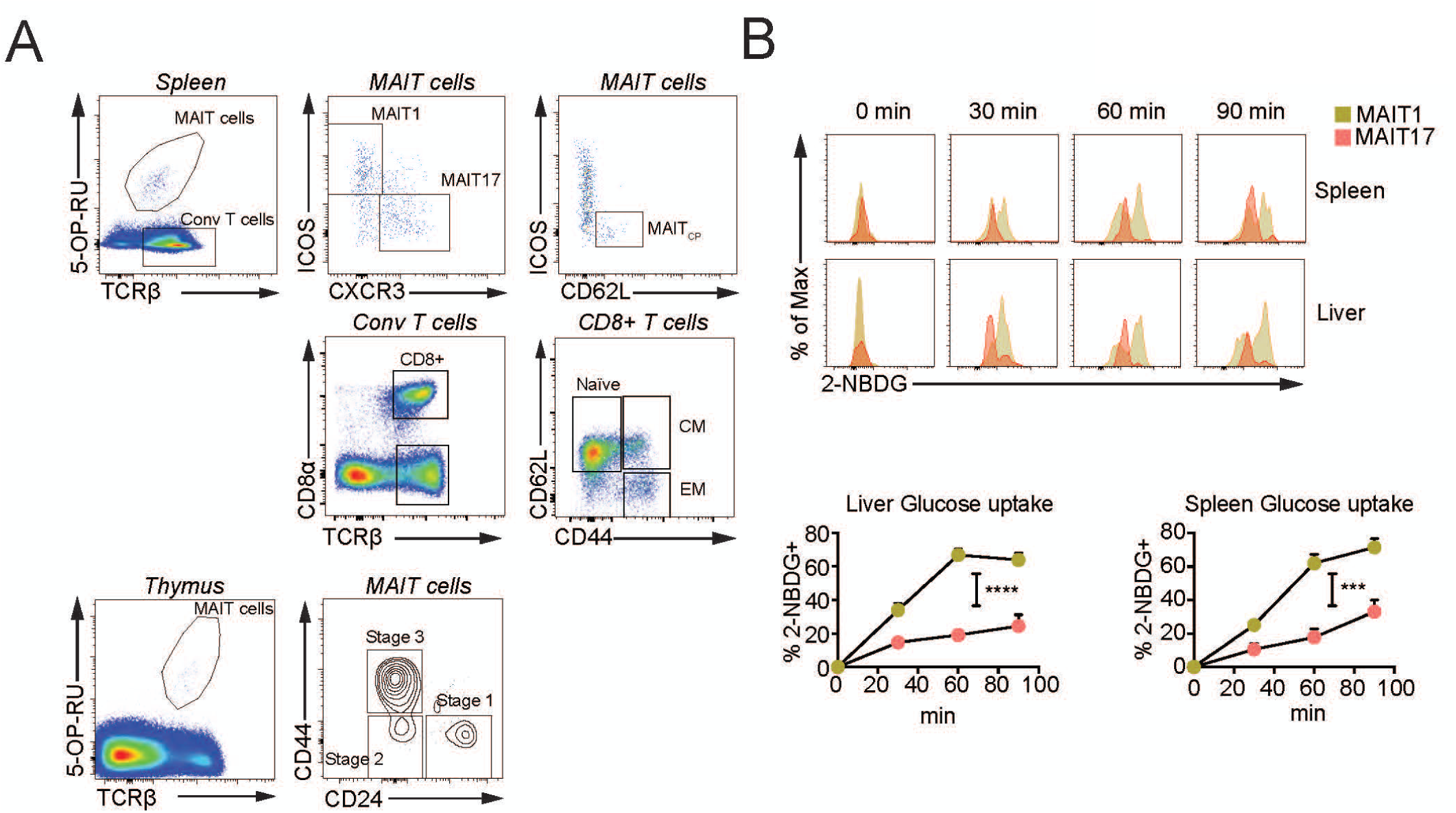
Mouse MAIT17 cells have high metabolic activity. (A) Gating strategy for subsets of MAIT and conventional (conv) or mainstream CD8^+^ T cells. MAIT cells were divided into MAIT17 and MAIT1 subsets based on ICOS and CXCR3 expression. Thymus MAIT cells were divided into various stages based on CD24 and CD44 expression. Spleen TCRβ^+^ CD8α^+^ T cells, excluding MAIT cells, were subdivided into naïve, central memory (CM) and effector memory (EM) subsets based on expression of CD62L and CD44. (B) Cells were isolated from indicated tissues and kinetics of fluorescent glucose (2-NBDG) uptake in MAIT1 (yellow) and MAIT17 (red) cell subsets was quantified; representative histograms (left) and quantification (right). Timepoints represent technical replicates from 8 pooled mice. Data analyzed by 2-way ANOVA with Geisser-Greenhouse correction, displayed as mean±□□SEM, \**P*□<0.05, \*\**P*□<0.01 \*\*\**P*□<0.001 and \*\*\*\**P*□<0.0001.

**Extended Data Figure 6:**
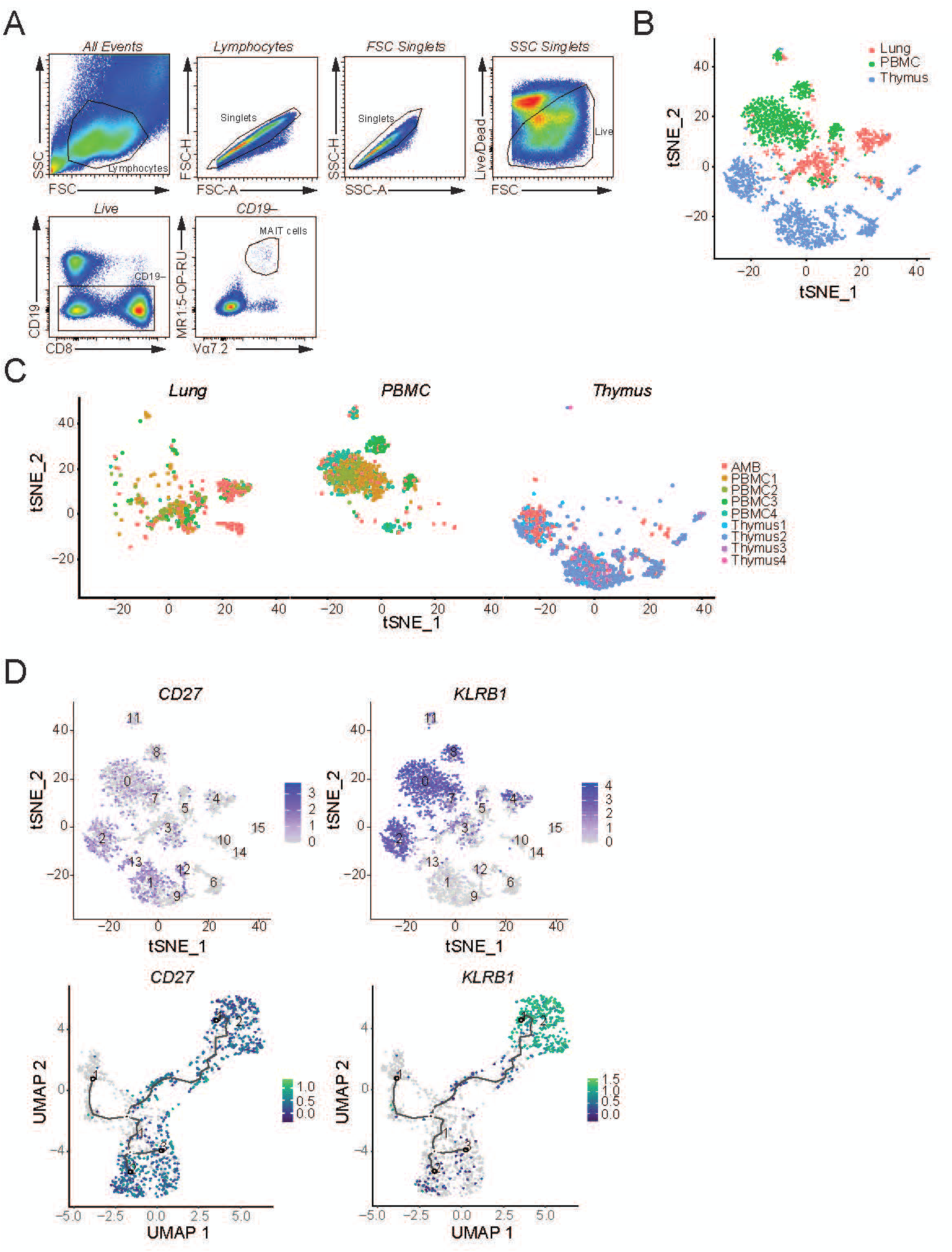
Human tissues have a distinct transcriptional signature. (A) Representative flow cytometry data used to identify MAIT cells in different human tissues. Live/Dead Yellow negative single cell events were gated by excluding B cells (CD19^+^). MAIT cells were identified as Vα7.2 TCR^+^ and 5-OP-RU human MR1 tetramer^+^ cells. (B) t-SNE plots representing human MAIT cells colored by their origin from different sites. (C) t-SNE representing cells from different donors (numbered) split by indicated tissues after de-multiplexing. AMB (Ambiguous) represents cells with no donor assignment. (D) t-SNE showing expression of *CD27 and KLRB1* transcripts in all human MAIT cells (upper plots) and expression of *CD27* and *KLRB1* along the pseudotime trajectory for human MAIT thymus cells as constructed by Monocle 3 (lower plots).

**Extended Data Figure 7:**
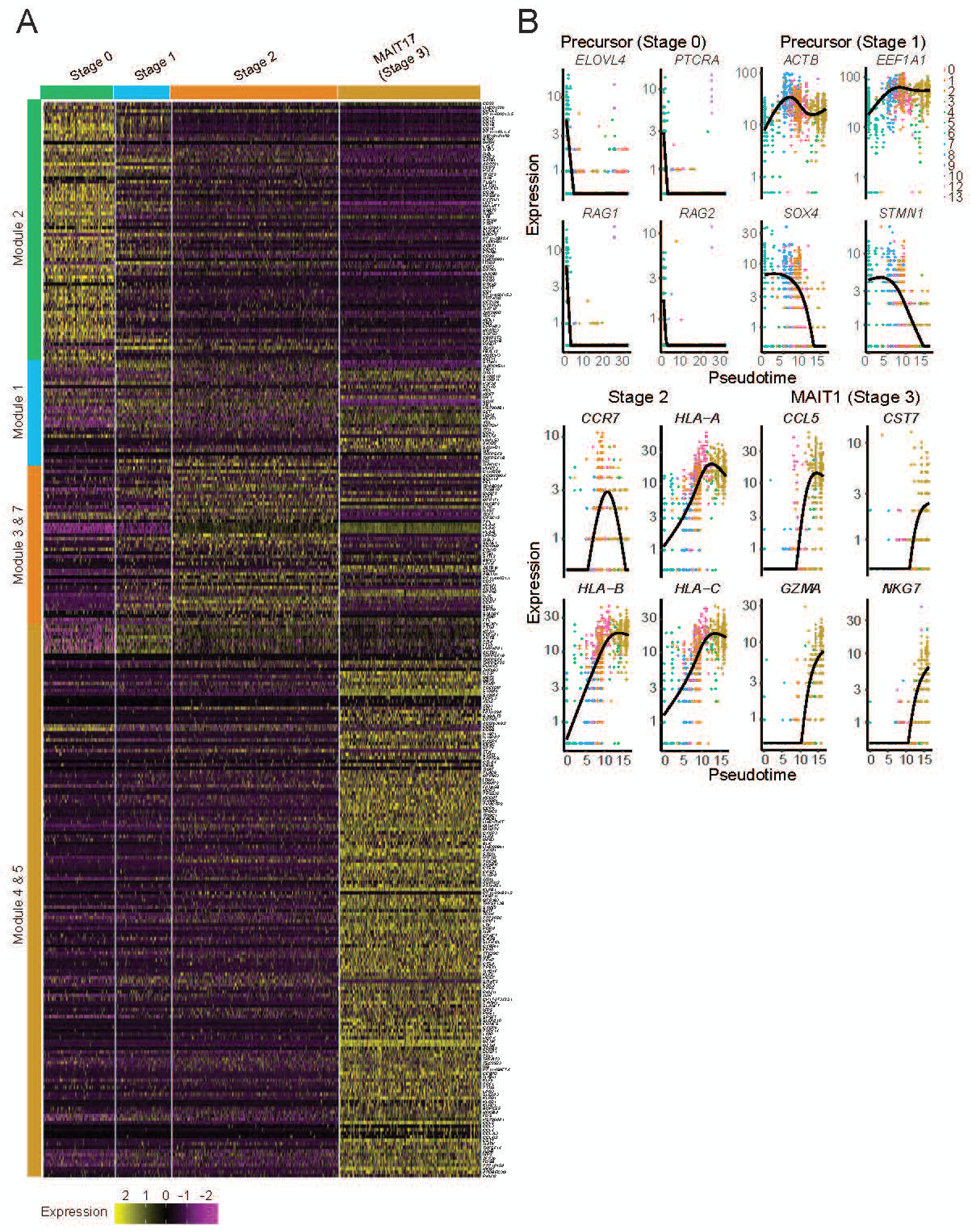
Transcriptional signatures reflect different stages of MAIT cell development in human thymus. (A) Scaled average expression heatmap of all the significantly differentially expressed genes along the MAIT cell thymus trajectory with Morans_I >0.2. Heatmap shows cells from the indicated clusters and MAIT cell differentiation stage on the top row. The gene modules to which the stage-specific genes belong are shown on the *y*-axis. (B) Expression of the indicated stage-specific genes along the pseudotime trajectory as constructed by Monocle 3.

**Extended Data Figure 8:**
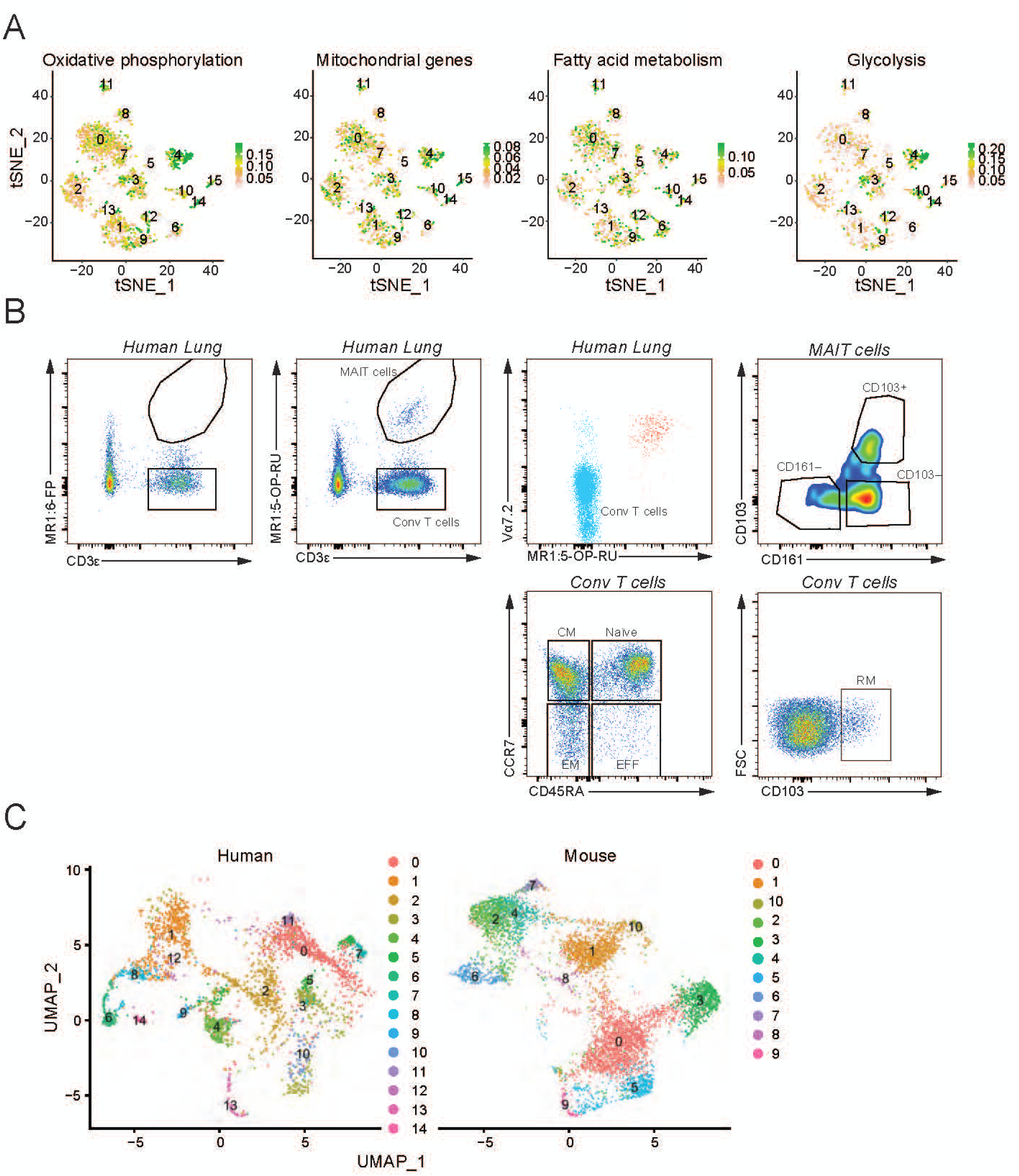
Metabolic signature of human MAIT cells and integration of human and mouse dataset. (A) t-SNE showing the oxidative phosphorylation, mitochondrial genes, fatty acid metabolism and glycolysis signature scores for each human MAIT cell. (B) Cells were isolated from human lung biopsies and stained with non-specific tetramer control (6FP, top) or 5-OP-RU loaded MR1 tetramer to identify MAIT cells (bottom). CD3^+^ 5-OP-RU-tetramer^+^ MAIT cells were further tested for Vα7.2 TCR alpha chain positivity and subdivided into three subsets based on expression of CD161 and CD103, as shown. TCRβ^+^ CD8^+^ T cells excluding MAIT cells were subdivided into naïve, central memory (CM), effector memory (EM) and resident memory (RM) subsets based on expression of CD45RA, CCR7 and CD103. (C) Integrated UMAP split by mouse and human showing clusters of cells with same coordinates as in the Fig 7A. Cells are labelled according to the cluster numbers in Fig 5 A (human) and Fig 1A (mouse).

**Extended Data Figure 9:**
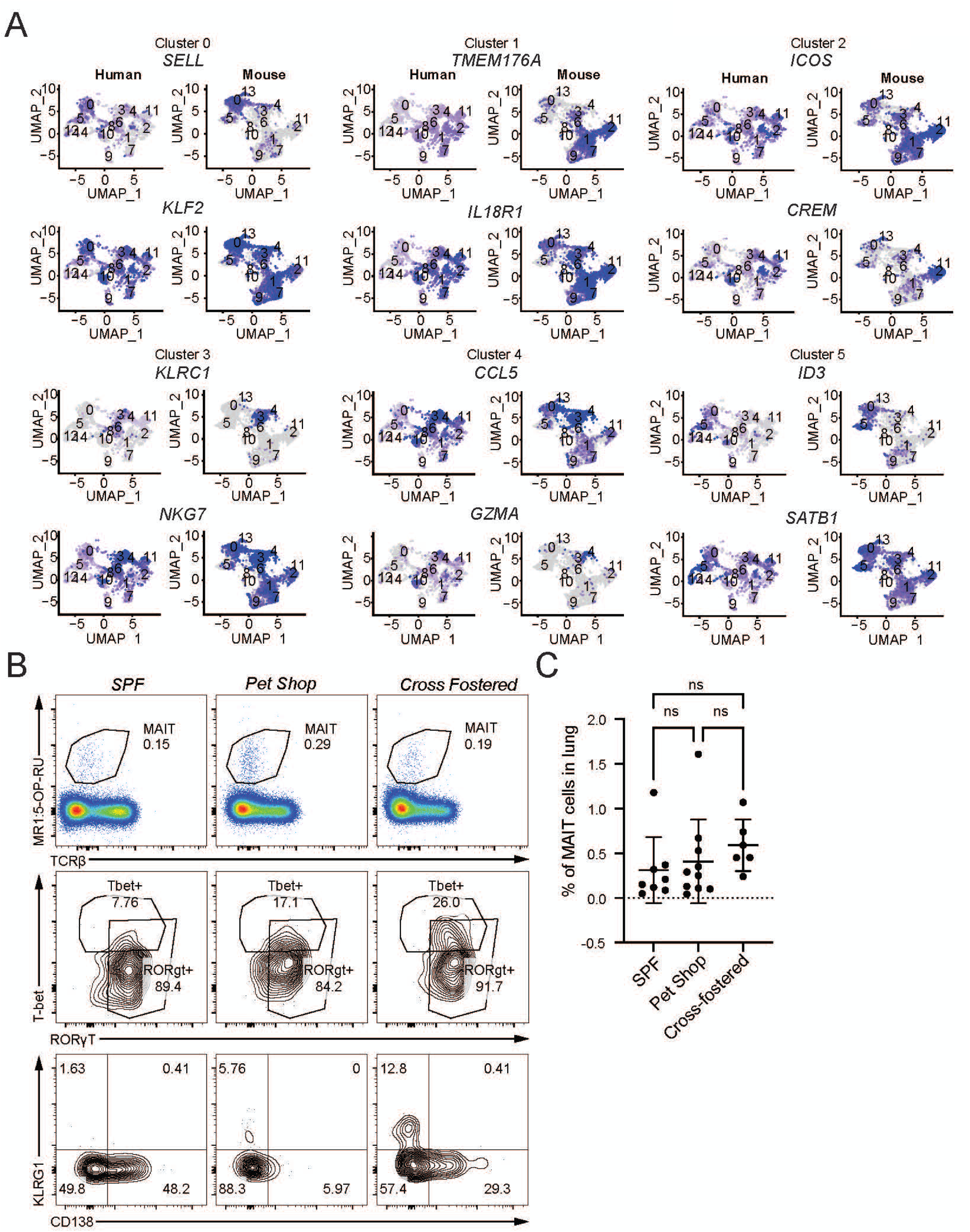
Comparison of human and mouse transcriptome and analysis of MAIT cells in pet shop and cross-fostered mice. (A) UMAPs showing expression by human and mouse MAIT cells of the top two marker genes of the indicated *i*-clusters. (B) Representative FACS plots from SPF, pet store and cross-fostered mice showing percentage of MAIT cells, intracellular expression of transcription factors T-bet and RORγT and surface expression of KLRG1 and CD138. (C) Percentage of MAIT cells in SPF, pet store and cross-fostered mice in lung. Data analyzed by one-way ANOVA with Tukey test displayed as mean± S.D. SPF mice n = 8, Pet shop mice n=10 and Cross-fostered mice n=6.

